# A unified model for the dynamics of ATP-independent ultrafast contraction

**DOI:** 10.1101/2022.10.14.512304

**Authors:** Carlos Floyd, Arthur T. Molines, Xiangting Lei, Jerry E. Honts, Fred Chang, Mary Williard Elting, Suriyanarayanan Vaikuntanathan, Aaron R. Dinner, M. Saad Bhamla

**Affiliations:** Department of Chemistry and James Franck Institute, University of Chicago, Chicago, IL 60637; Department of Cell and Tissue Biology, University of California, San Francisco, CA 94143; School of Chemical and Biomolecular Engineering, Georgia Institute of Technology, Atlanta, GA 30318; Department of Biology, Drake University, Des Moines, IA 50311; Department of Physics, North Carolina State University, Raleigh, NC 27607

**Author notes:** Electronic mail (Mary Williard Elting). Electronic mail (Suriyanarayanan Vaikuntanathan). Electronic mail (Aaron R. Dinner). Electronic mail (M. Saad Bhamla).

## Abstract

In nature, several ciliated protists possess the remarkable ability to execute ultrafast motions using protein assemblies called myonemes, which contract in response to Ca^2+^ ions. Existing theories, such as actomyosin contractility and macroscopic biomechanical latches, do not adequately describe these systems, necessitating new models to understand their mechanisms. In this study, we image and quantitatively analyze the contractile kinematics observed in two ciliated protists (*Vorticella sp* and *Spirostomum sp*), and, based on the mechanochemistry of these organisms, we propose a minimal mathematical model that reproduces our observations as well as those published previously. Analyzing the model reveals three distinct dynamic regimes, differentiated by the rate of chemical driving and the importance of inertia. We characterize their unique scaling behaviors and kinematic signatures. Besides providing insights into Ca^2+^-powered myoneme contraction in protists, our work may also inform the rational design of ultrafast bioengineered systems such as active synthetic cells.

## I. INTRODUCTION

A well-known molecular mechanism for motion in biological systems is actomyosin contractility, which involves the relative sliding of actin filaments due to ATP hydrolysis by bound molecular motors^1^. However, the fastest motions currently known to occur in the biological world (measured as the proportional change in length per unit time) are due not to actomyosin assemblies but to lesser-known protein assemblies called myonemes^2–4^. We use “myonemes” in the broadest sense^5^, encompassing terms used for specific organisms such as “spasmonemes” and “M-bands.” Myonemes are common in ciliated protists: for example, they are found in the wine glass-shaped genus *Vorticella*^6,7^, the cigar-shaped genus *Spirostomum*^8–11^, the trumpet-shaped genus *Stentor* ^12,13^, and the chandelier-shaped genus *Zoothamnium*^14,15^. Similar Ca^2+^-responsive supramolecular assemblies are proposed to occur in a range of other single-celled organisms^16–23^, and possibly even plants^24–26^. The mechanism of ultrafast myoneme contraction is qualitatively different from actomyosin contraction in that ATP is not directly consumed during the contractile motion^6,27^. Instead, a local increase in Ca^2+^ concentration acts as a mechanical actuator to initiate contraction^28–31^. ATP is subsequently consumed to pump Ca^2+^ away from the myoneme fibers and reset the process^14,32^.

While several experimental studies have been conducted on myoneme-based contraction dynamics^7**–**10,^, it is difficult to draw general conclusions without a theoretical framework to interpret the observations. Whereas actomyosin contractility has been extensively studied theoretically and computationally^42–47^, Ca^2+^-powered myoneme-based contractility is relatively unexplored from a modeling point of view. An existing computational model of *Vorticella*^48^ is specific to the geometry of the organism and thus does not apply across species. Previous mechanochemical analyses also do not address the dynamics of contraction^6,9^. Questions that a mathematical model of myoneme contraction might address are: what synchronizes contraction across the organism’s length, what sets the rate of chemical driving, and what are the key geometric and mechanical factors that affect the contraction dynamics?

Here, we develop a minimal mathematical model of myoneme contraction and show that it applies across species (Figure 1). In our model, an elastic medium in a dissipative environment is driven out of equilibrium by a traveling wave of shrinking rest lengths, which captures the effect of a traveling Ca^2+^ wave. This simple setup omits several system-specific details but can fit experimental contraction measurements for *Spirostomum* and *Vorticella*. Mathematically, the model comprises a wave equation with dissipation and a non-autonomous source term. We derive an approximate form for the non-autonomous function by analyzing an auxiliary kinetic model for a traveling Ca^2+^ wave. Using the model, we show that the balance between chemical driving, mechanical stiffness, and viscous drag results in three dynamic regimes. These regimes are distinguished by the relative importance of inertia and Ca^2+^ wave speed, and they exhibit kinematic signatures, including asymmetric contraction and different scaling behavior of the maximum contraction rate. This work provides a starting point for extracting general mechanochemical principles of myoneme contraction in biological and engineered systems.

**FIG. 1.**
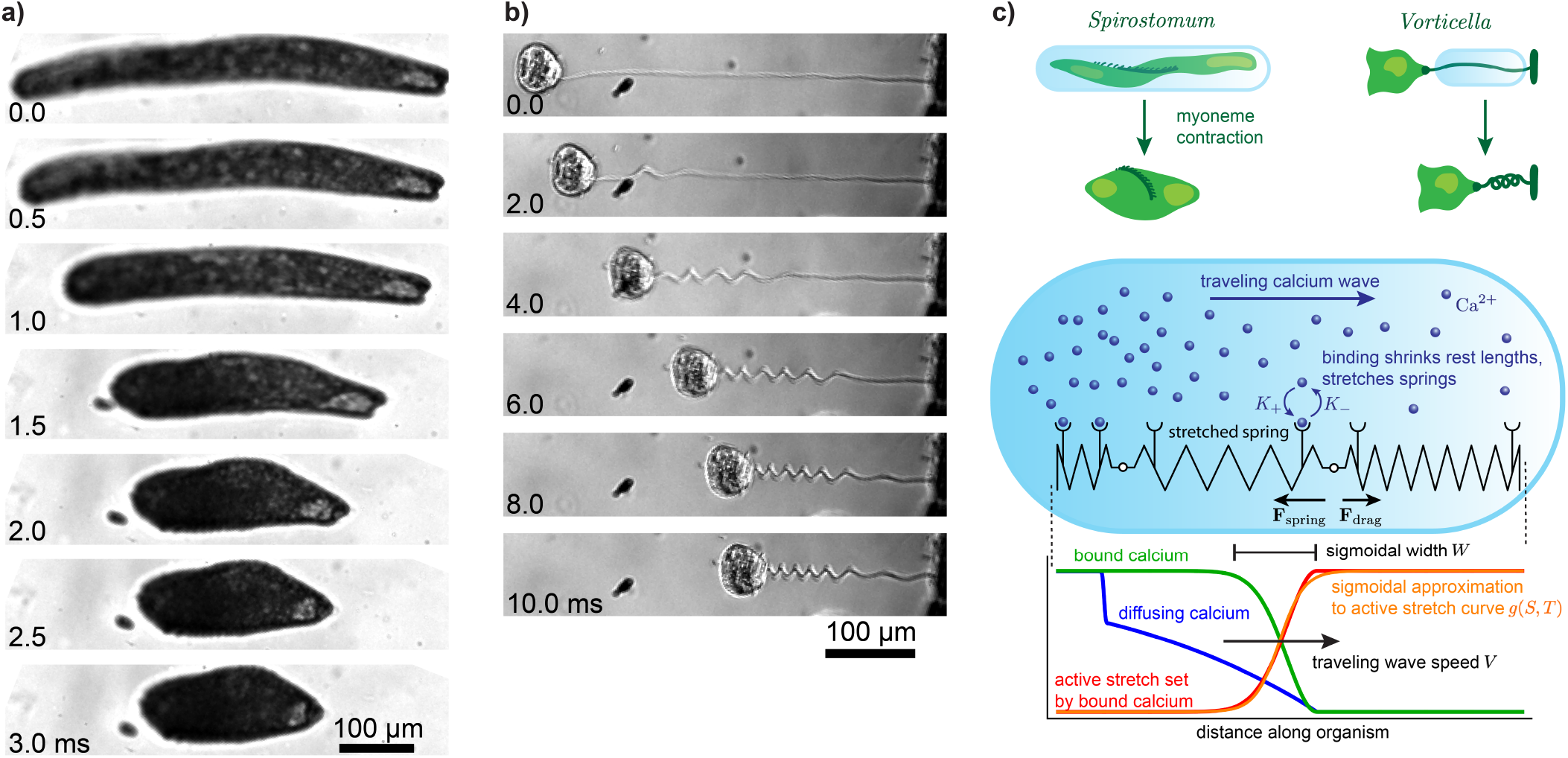
Myoneme-based contraction in ciliated protists. (a) Images of *Spirostomum* undergoing contraction, with the time in milliseconds labeled on the bottom left of each image. Rapid contraction begins at *T* = 0 ms and is followed by a slower re-elongation phase (not depicted). Note that the left side of the organism begins contracting (between 0.5 and 1.0 ms) before the right (between 1.0 and 1.5 ms). (b) Same as (a) for *Vorticella*. (c) Schematic of the model. The contractile body of *Spirostomum* and stalk of *Vorticella* are abstracted as a 1D elastic medium in a viscous environment, represented as a chain of springs. The local spring rest length (depicted as the number of ridges) depends on the number of bound Ca^2+^ ions (shown as blue spheres bound to receptors on the springs). A rightward traveling Ca^2+^ wave thus shrinks the spring rest lengths, exerting forces in the chain which then contracts against viscous drag. The bottom graph schematically illustrates several quantities in our mechanochemical model at a fixed time, which we elaborate on in the main text.

## II. MODEL DERIVATION

Myoneme contraction is driven by an increase in cytosolic Ca^2+^, which occurs as a traveling wave that traverses the organism’s length in several milliseconds^29,48–51^. The mechanism by which Ca^2+^ physically induces contraction is believed to be a screened entropic spring effect: Ca^2+^ binds to the negatively charged myoneme fibers and screens their electrostatic self-repulsion, which allows for the entropic contribution to the free energy to induce collapse of the polymer^52^. This system is sometimes considered a polyelectrolyte gel, in which thermodynamic balances of entropic and osmotic forces determine the gel’s equilibrium^14,53,54^. The precise contributions to the myoneme free energy as a function of length are not needed to formulate a minimal mechanical model, and we make the phenomenological assumption that the system obeys linear elasticity^55^. The key difference from the usual elastic theory is that the rest lengths of the elastic elements are dynamic quantities that depend on the local concentration of bound Ca^2+^ (Figure 1).

To formulate the model mathematically we start from a discrete chain of harmonic springs in one dimension with externally imposed rest lengths, and we define *X*_*i*_(*T*) as the position of the *i*^th^ mass at time *T*. The spring stiffness parameter *k*, mass *m*, drag *γ*, and initial rest length *δ*, are the same for all springs, and the instantaneous rest length 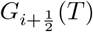 of the spring connecting masses *i* and *i* + 1 is a non-autonomous function which we specify later. The force on mass *i* is

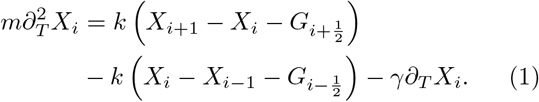

Rather than a discrete description in terms of the index *i*, we seek a continuum description of the dynamics in terms of a material coordinate *S* ∈[0, *L*], where *L* is the uncontracted length of the system. By Taylor expanding the functions *X* and *g* ≡*G/δ* to second order in *δ* and then letting *δ* →0, we find the following partial differential equation (PDE) for the position *X*(*S, T*):

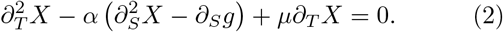

Here, 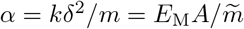 is the product of the Young’s modulus *E*_M_ and the effective spring crosssectional area *A* divided by the linear mass density 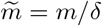 (where we used *k* = *E*_M_*A/δ*^55^). The parameter *α* can be interpreted as the square of the speed of sound in the spring chain^56^ 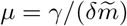 is the ratio of the drag constant per unit length to the linear mass density, and *g*(*S, T*) is a non-autonomous function for the active stretch, defined as the ratio of the current spring rest length *G*(*S, T*) to the initial rest length *δ*. The quantities *g, α*, and *μ* remain finite as *δ* →0 because *G* ∼*δ, k* ∼*δ*^*−*1^, *m* ∼*δ* and *γ*∼ *δ*, while *E*_M_, *A*, and 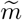 are independent of *δ*. Eq. (2) represents the force balance of *X*(*S, T*), where the first term is the acceleration, the second term (proportional to *α*) captures the force due to the difference between the local stretch and the active stretch *g*, and the third term captures the drag force. See *SI Appendix Supplementary methods* Section B 1 for a detailed derivation of the model.

We find the boundary conditions by retaking the continuum limit at the terminal masses in the discrete representation (Eq. (1)), noting that these masses are connected to only a single spring. This yields the stress-free condition

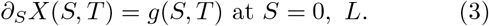

In the case of *Vorticella*, which has one end (say *S* = *L*) attached to a fixed substrate, we use the boundary condition *X*(*L, T*) = *L* at that end. We further take the initial conditions

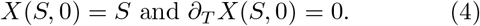

We model the active stretch *g*(*S, T*) as a linear function of the local concentration of Ca^2+^ bound to the myoneme fibers; see Eq. (17). Specifying the active stretch function *g*(*S, T*) requires describing the Ca^2+^ dynamics, for which we adapt a tractable reaction-diffusion model of a traveling Ca^2+^ wave^57^, as described in *Materials and methods* Section V A and See *SI Appendix Supplementary methods* Section B 2. By analyzing this model, we find that a good approximation to the active stretch function *g*(*S, T*) = *g*^*\**^(*S* − *V T* ; *W*) is a traveling wave moving at speed *V* and having a sigmoidal profile *g*^*\**^ with width *W* (Eq. (22); Figure 1c). This function decreases from 1 to *g*_min_ ∈[0, 1] as the wave passes through the organism. An additional parameter *O* sets the time offset such that the center of the wave arrives at *S* = 0 at time *O*. As described in *Results* Section III B, we can approximately express *V* and *W* in terms of the kinetic parameters of the Ca^2+^ model (Eq. (5) and Eq. (B31)).

Our model neglects contributions from transverse dimensions and twisting effects, which may alter the onedimensional dynamics through elastic coupling in the real system^58,59^. As explained in *SI Appendix Supplementary methods* Section B 3, we assume that this coupling is negligible as a first approximation. In that section, we also discuss the explicit assumptions needed to disregard the poroelastic contribution arising from fluid motion around the organelles inside the contracting organism. We demonstrate that under standard approximations made when treating cytoplasm poroelasticity^60^, poroelasticity in our model contributes an additive term to the Stokes drag coefficient (see *SI Appendix Supplementary methods* Eq. (B48)). Moreover, we examine a model extension in which motion encounters resistance from auxiliary elastic elements with unchanged rest lengths during Ca^2+^ binding. This representation approximates, for instance, the microtubule sheath under the cortex of *Spirostomum*^11,35^ or the elastic stalk supporting the thin spasmoneme fiber in *Vorticella*^48^. We discover that a linear change of variables effectively maps the model with auxiliary elastic elements to the model presented here.

## III. RESULTS

### A. A minimal continuum model reproduces contraction kinematics

Despite the model’s simplicity, we find that it reproduces the contraction kinematics of *Spirostomum* and *Vorticella* semi-quantitatively. We illustrate this by comparing the trajectories of normalized length *λ*(*T*) ≡ (*X*(*L, T*) −*X*(0, *T*)) */L* from experimental measurements to fitted model curves. Contraction events for *Spirostomum* and *Vorticella* are shown in *SI Appendix Supplementary movies* S1 and S2, and the experimental methods are described in *Materials and methods* Section V B. The second-order non-autonomous PDE model does not have an exact closed form solution, though we describe exact series solutions in two limiting cases in *Results* Section III C. To fit the model, we numerically integrate the PDE with different choices of parameters and compare the results to the experimental traces. See *Materials and methods* Section V C for details on the computational methods. In Figure 2a, we show five contraction measurements in *Spirostomum* along with a model solution fitted to their average, illustrating the feasibility of our model in capturing the contraction dynamics. The corresponding contraction velocities |*∂*_*T*_ *λ*| are shown in Figure 2b. We also fit the model with a fixed boundary condition at *S* = *L* to data for five *Vorticella* contraction events (Figures 2c and d).

**FIG. 2.**
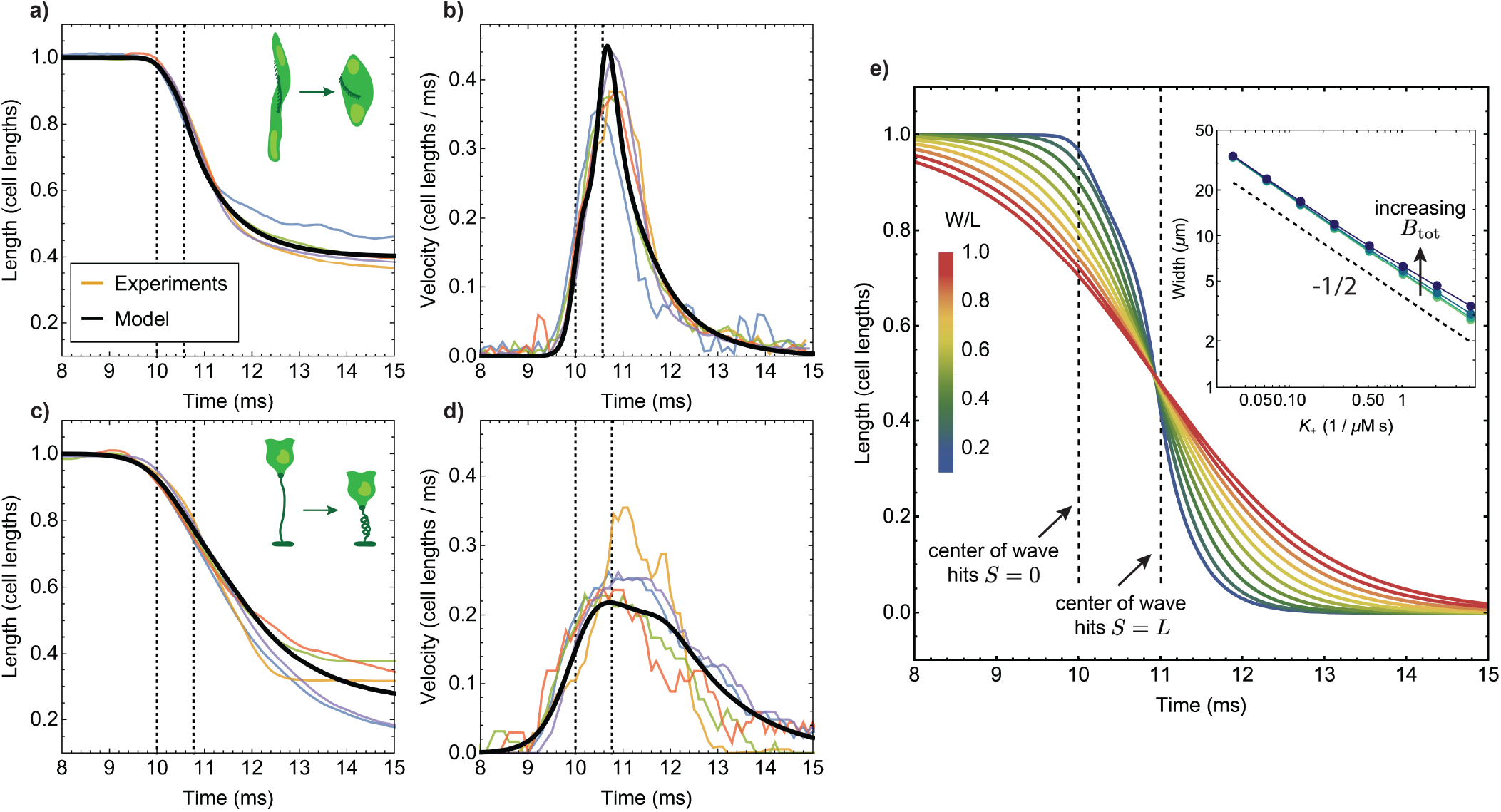
Comparison of the model to experimental data. (a) Five measurements of *Spirostomum* contraction, collapsed to the same scale by normalizing length using the organism’s uncontracted length *L* and by offsetting time so that contraction begins around *T* = 10 ms. Thin lines represent individual contraction events, and we color each differently. The thick black curve shows the model fit to all of these trajectories at once; see *SI Appendix Supplementary results* Section A 2 for fits to the individual curves. Vertical dashed lines indicate the times in the model solution when the center of the active stretch wave arrives at *S* = 0 and *S* = *L*. Time is shifted by choosing *O* = 10 ms in Eq. (22) to achieve numerical agreement between the boundary and initial conditions (see *Material and methods* Section V C). (b) The corresponding rate trajectories, obtained by numerically differentiating the data in (a). Colors of individual events are the same as in (a). (c) and (d) The same as (a) and (b) but for *Vorticella* contraction. (e) Simulated length trajectories with *V* = 1 *L/*ms, *α* = 2 (*L/*ms)^2^, *g*_min_ = 0, *O* = 10 ms, and *μ* = 10 ms^*−*1^. The width *W* of the wave is varied from 0.1 *L* to 1.0 *L* in steps of 0.1 *L* as the colors range from blue to red. The inset shows how *W* depends on the binding rate *K*_+_ in the Ca^2+^ model. The total myoneme concentration *B*_tot_ ranges from 500 to 4, 000 *μ*M in multiples of 2 as the colors range from light green to purple. The dashed line in the inset represents the analytically predicted scaling 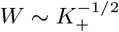. For the remaining parameters used in the inset, see *SI Appendix Supplementary methods* Table IV.

In *SI Appendix* Figures 12 and 13, we present fits for each of the ten individual trajectories shown in Figures 2a and c, demonstrating that we can obtain excellent fits for all trajectories using fairly consistent parameters for each organism. As we explain in the Results Section III C, we find that these contraction events occur in a regime where inertia is unimportant, meaning that the dynamics are approximately determined only by the ratio *μ/α*. We fit the simplified “slow-wave” model (cf. Eq. (10)) and obtain a typical set of parameters for *Spirostomum*: *L* = 0.58 mm, *g*_min_ = 0.35, *μ/α* = 5 ms*/*mm^2^, *V* = 0.8 mm*/*ms, and *W* = 0.18 mm. For *Vorticella*, a typical set of parameters is *L* = 0.27 mm, *g*_min_ = 0.13, *μ/α* = 2 ms*/*mm^2^, *V* = 0.15 mm*/*ms, and *W* = 0.06 mm. In *SI Appendix Supplementary results* Section A 2, we demonstrate that the fitted parameters for *Vorticella* are consistent in order of magnitude with physiological parameters used in a detailed computational model in Ref. 48. According to a two-tailed *t*-test, the distributions of quantities *μ/α, V*, and *W* each have means that are statistically different for the two organisms (*t*-test *p*-value ≪0.05). In the following section, we investigate the kinetic factors that underlie the rate of chemical driving captured by the parameters *V* and *W*.

### B. Ca^2+^ dynamics control contraction

The speed *V* and, less intuitively, the width *W* of the active stretch wave *g*(*S* −*V T* ; *W*) determine how rapidly the length changes over time. As *W* decreases, the active stretch wave approaches a traveling step function in which the springs ahead of the wave have their original rest lengths, and those behind have rest lengths *g*_min_. As *W* increases, the distribution of changing rest lengths widens, and the driving and relaxation become more spatially and temporally homogeneous. We illustrate this in Figure 2e, where we show the length trajectory for several values of *W*.

We are interested in understanding which features of the Ca^2+^ dynamics control the quantities *V* and *W*, since these parameters control the rate at which the mechanical system is chemically driven out of equilibrium. To do this, we analyzed a kinetic model^57^ for a traveling Ca^2+^ wave powered by calcium-induced calcium release^61,62^ from an out-of-equilibrium internal store (e.g., the endoplasmic reticulum^29,49^) into the cytosol. It has been previously shown that in this model, the wave speed *V* approximately follows^57^

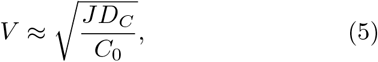

where *J* represents the Ca^2+^ influx of open channels, *D*_*C*_ is the diffusion constant of Ca^2+^, and *C*_0_ is the concentration threshold of the Ca^2+^ channels (see *Materials and methods* Section V A and *SI Appendix Supplementary methods* B 2 for more details). We incorporated the binding of cytosolic Ca^2+^ to myoneme fibers into the model and derived a new relationship for the width *W* of the resulting active stretch wave:

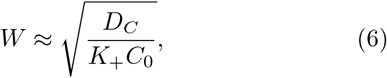

where *K*_+_ is the rate at which Ca^2+^ binds to myonemes. We provide numerical confirmation of this relationship in *SI Appendix Supplementary methods* Section B 2 and the inset of Figure 2e. A seemingly restrictive assumption made in the derivation of Eq. (B31) is that the total concentration of myoneme binding sites *B*_tot_ is much smaller than the typical cytosolic Ca^2+^ concentration. However, we find numerically that the predicted scaling holds even as *B*_tot_ increases to relatively large values.

This analysis reveals that features in the dynamics of contraction, such as the time taken for the length to change, can inform us about the magnitudes of the underlying biophysical parameters like the binding rate *K*_+_. Assuming *D*_*C*_ remains the same in both organisms (though the presence of internal buffers could affect this^57^), the estimated values of *V* and *W* for *Vorticella* and *Spirostomum* suggest, through Eqs. (5) and (B31), that these organisms possess different threshold Ca^2+^ concentrations *C*_0_^29,63,64^ and different release rates *J* or binding constants *K*_+_. This minimal model serves as a first step in connecting contraction dynamics to the organisms’ biochemistry, but more *in vivo* experiments are necessary to test these predictions.

### C. Dimensional analysis reveals the minimal set of model parameters

The model considered so far (Eq. (2) together with Eq. (22)) is written in dimensional quantities and depends on the parameters *α* and *μ*, which are related to the stiffness of the material and viscosity of the environment, and *V, W*, and *g*_min_, which are the speed, width, and minimum value of the active stretch wave. However, these parameters are not all independent. For example, rescaling time or space by a common factor would leave the dynamics unchanged. We now show that it is possible to reduce the number of parameters by one through nondimensionalization. We use lowercase variables throughout to denote dimensionless quantities.

There are two length scales, *W* and *L*, and we pick the uncontracted length *L* as the reference length and define *s* = *S/L, x* = *X/L*, and *w* = *W/L*. Next, three timescales can be defined:

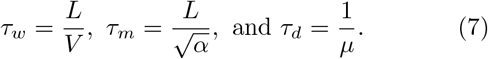

Here, *τ*_*w*_ is the wave timescale, which represents the time taken for the center of the traveling wave to traverse the length *L*; *τ*_*m*_ is the material timescale, which represents the time taken for a sound wave to traverse the length of the spring chain; *τ*_*d*_ is the drag timescale, characterizing the exponential deceleration of motion due to viscous damping. Unless otherwise specified, we non-dimensionalize time using *t* = *T/τ*_*w*_. In these units, the wave speed is equal to 1, and Eq. (2) is expressed as the following PDE for *x*(*s, t*):

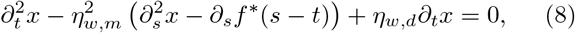

where *f* ^*\**^(*s*) ≡*g*^*\**^(*Ls*) is the active stretch wave profile, rewritten to take a dimensionless argument *s*. The size of *τ*_*w*_ relative to the two remaining timescales defines the non-dimensional parameters

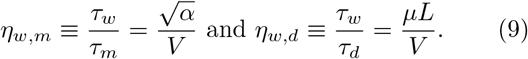

The parameter *η*_*w,m*_ describes the ratio of the myonemal speed of sound (which depends on the stiffness and density) to the wave traversal speed, and it is independent of length. The parameter *η*_*w,d*_ describes the ratio of the drag timescale (depending on viscosity and density) to the wave traversal speed and is proportional to the length.

We can further simplify these non-dimensional dynamics by taking the “slow wave” limit *η*_*w,m*_, *η*_*w,d*_ ≫ 1, in which Eq. (8) reduces to

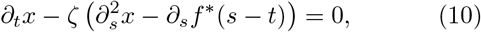

where the only remaining parameter is

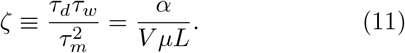

Eq. (10) can be viewed as an over-damped limit of the model, in which inertia has dropped out of the dynamics because only a first derivative with respect to time remains, and in which mass is no longer relevant because *ζ* is independent of 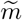.

Eq. (10) is similar to the diffusion (or heat) equation with a source term. One difference is the boundary conditions: typically, the heat equation has fixed Dirichlet or Neumann boundary conditions, yet *x* obeys a boundary condition that depends in time on *f* ^*\**^(*s* − *t*) (cf. Eq. (3)).

However, we can define

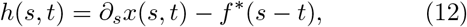

and express Eq. (10) as

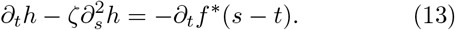

The boundary conditions are now the fixed Dirichlet conditions *h*(0, *t*) = *h*(1, *t*) = 0, and thus Eq. (13) is exactly the diffusion equation with a source term −*∂*_*t*_*f*, for which exact series solutions can be found. The mixed boundary conditions of *Vorticella* also result in mixed boundary conditions for the diffusion equation.

We also consider a “quench” limit *η*_*w,d*_, *η*_*m,d*_ *≪*1, in which the traveling wave moves at infinite speed, such that the rest lengths of all the springs are set to their new values before the organism can react. This implies that *f* (*s*) is independent of time. Because the timescale *τ*_*w*_ is zero if the wave is infinitely fast, we instead non-dimensionalize time using *τ*_*m*_ in the quench limit. The dynamics consist of only the subsequent relaxation of the system to its equilibrium

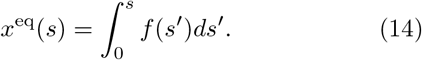

The deviation from equilibrium, *y*(*s, t*) ≡*x*(*s, t*) −*x*^eq^(*s*), obeys the dynamics

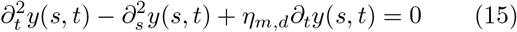

with the initial condition *y*(*s*, 0)) ≡*x*(*s*, 0) −*x*^eq^(*s*), *∂*_*t*_*y*(*s*, 0) = 0, and stress-free boundary conditions *∂*_*s*_*y*(0, *t*) = *∂*_*s*_*y*(1, *t*) = 0. We note that the one remaining parameter

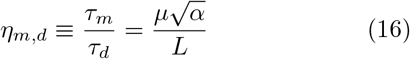

is independent of wave speed, which is infinite. The quench limit has an exact Fourier series solution, described in *SI Appendix Supplementary methods* Section B 4.

In Figures 3a and b we compare the solutions of the full non-dimensional model (Eq. (8)), the slow wave model (Eq. (10)), and the quench model (Eq. (15)) for two sets of the non-dimensional parameters *η*_*w,d*_ and *η*_*m,d*_. In these plots we fix *w* = 0.1 and *g*_min_ = 0 for simplicity. For *η*_*w,d*_ = *η*_*m,d*_ = 1 (Figure 3a), the models all behave quite differently because neither limiting case is reached. Decaying oscillations about *l* = 0 are observed for the full and quench model, while the slow wave model does not oscillate because it lacks inertia. The oscillations for the quench and full model have the same frequency (which is proportional to *η*_*m,d*_, reflecting the resonant frequency of the spring chain) yet different waveforms and amplitudes. For *η*_*w,d*_ = 20, *η*_*m,d*_ = 2 (Figure 3b), the full model and slow wave model are in close agreement.

**FIG. 3.**
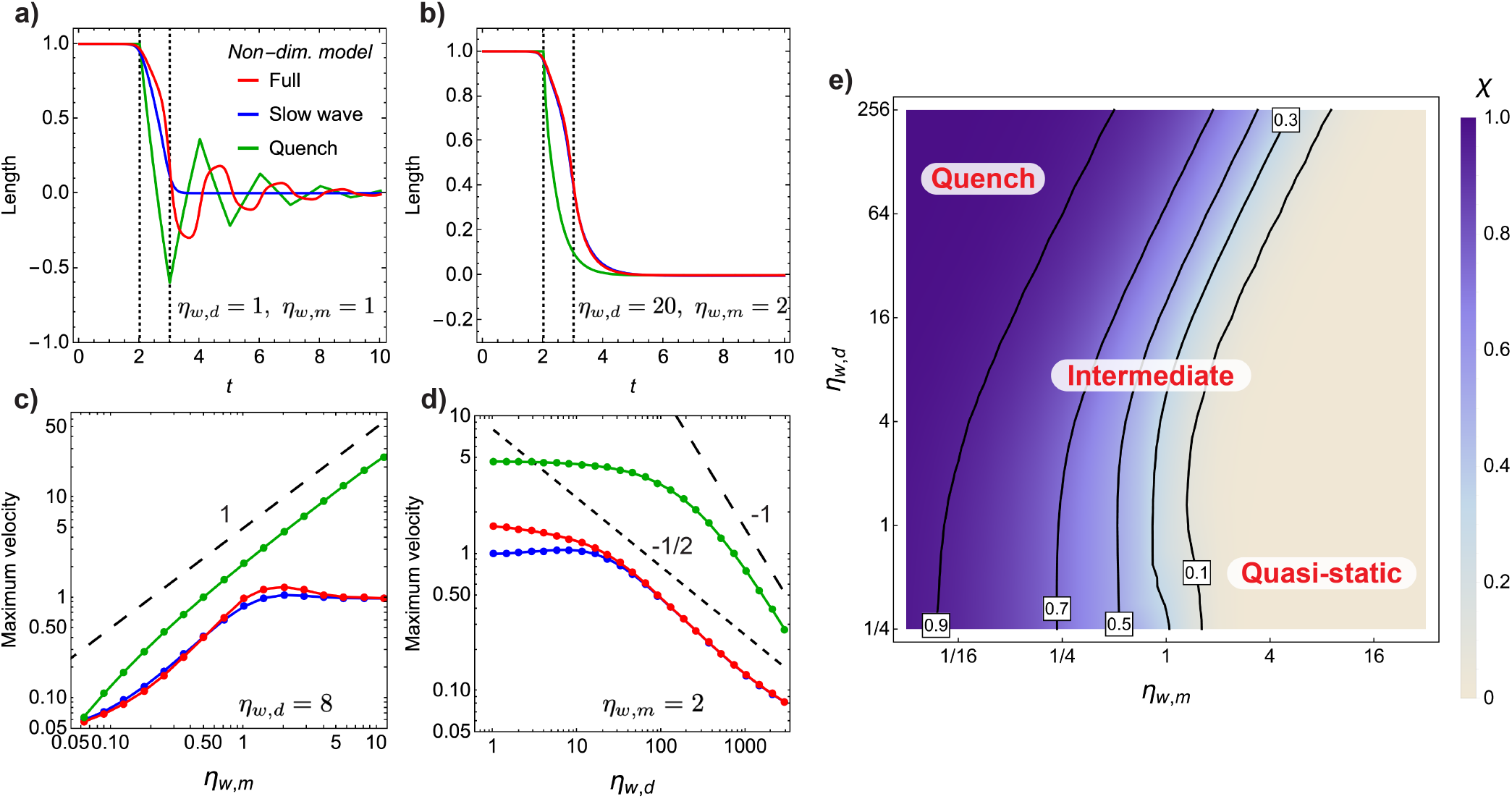
The non-dimensional model and its limiting versions capture key kinematic properties of contraction. (a) Non-dimensional length *λ*(*t*) as a function of time for the three versions of the non-dimensional model. Parameter values are *η*_*w,d*_ = *η*_*w,d*_ = 1, *w* = 0.1, *g*_min_ = 0. The vertical lines at *t* = 2 and 3 indicate the times when the center of the active stretch wave enters and exits the system. For the quench model, the quench occurs at *t* = 2. (b) Same as (a) but with *η*_*w,d*_ = 2 and *η*_*w,d*_ = 20. (c) and (d) Scaling of 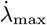 with *η*_*w,m*_ (c) and *η*_*w,d*_ (d) for the three models. The colors are the same as in (a). For panel (c), *η*_*w,d*_ is fixed at 8, while for panel (d), *η*_*w,m*_ is fixed at 2. (e) Heatmap of *χ*, the peak elastic energy stored in the system as a fraction of the maximum possible energy, as *η*_*w,m*_ and *η*_*w,d*_ are varied for the full model. Contours are drawn and labeled at selected values of *χ*.

A stark feature of the full model compared to the quench model is the asymmetry of contraction: in the quench model, all springs instantly have new rest lengths, so all springs relax simultaneously and symmetrically about the center *s* = 0.5. This is not the case for the full model, where the wave first arrives at *s* = 0, causing the springs to begin relaxing before those at *s* = 1. This asymmetry of contraction can be observed in the experimental images of *Spirostomum* and *Vorticella* in Figures 1a and b, and we further illustrate this in Figure 4a, b, and c. In Figure 4d, e, f, and g we show *x*(*s, t*) at several values of *t* for the full and quench models corresponding to the length trajectories shown in Figures 3a and b.

**FIG. 4.**
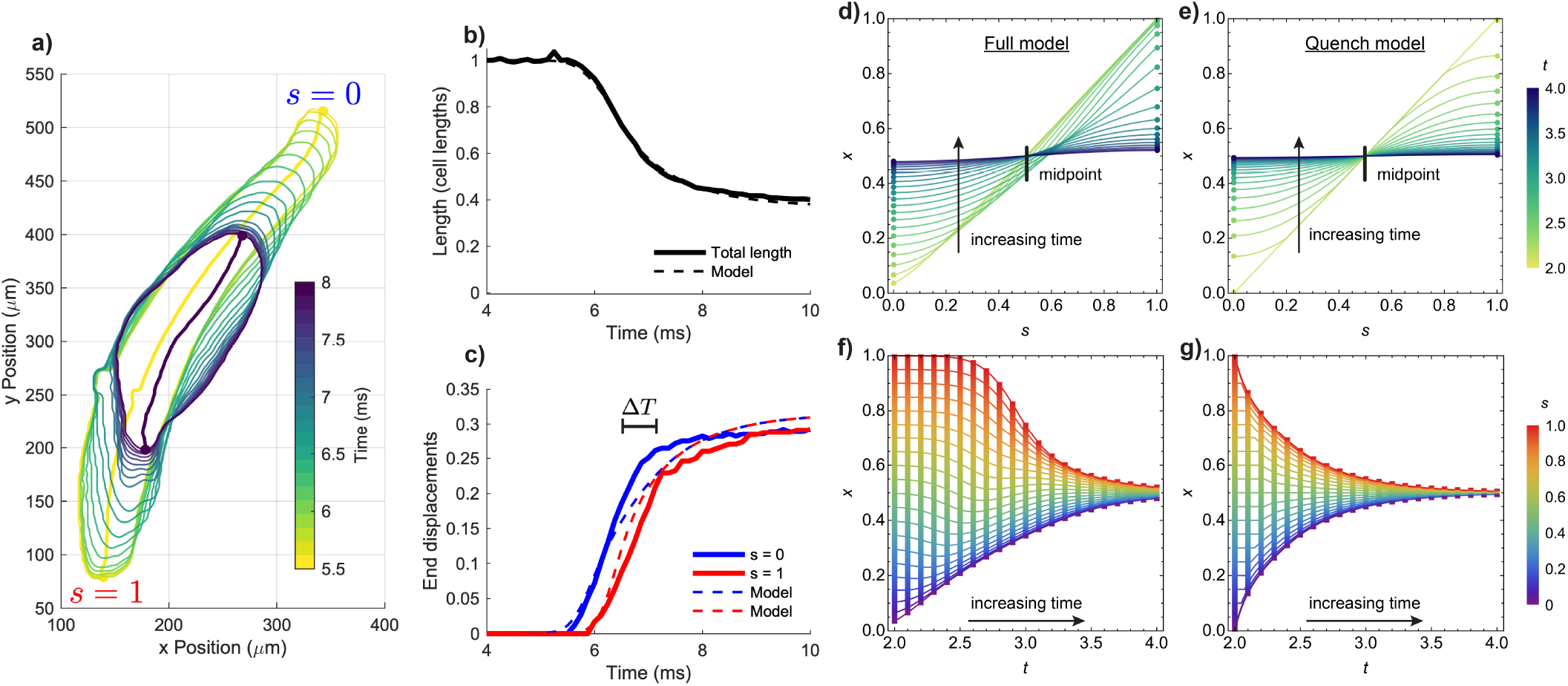
Asymmetric contraction of *Spirostomum*. (a) A sequence of contours of *Spirostomum* obtained from 8, 000 Hz video imaging. Colors range from yellow to purple as time ranges from 5.5 ms to 8 ms. The initial and final midlines, whose arc lengths determine the measured length of the organism, are drawn in yellow and purple. (b) The trajectory of the length during this contraction event. The dashed curved line shows the trajectory of the simulated model, which was fit to match the experimentally measured solid curve. (c) The displacement trajectories of the top (red) and bottom (blue) ends of the midlines and the corresponding quantities in the fitted model are shown as dashed curves. These displacements were measured by projecting the endpoints to the initial midline curve and computing the arc length from the end to the projected point. Asymmetric contraction can be observed as the top end begins contracting before the bottom end (by a time difference labeled Δ*T*), a feature that the fitted model reproduces. (d) and (e) Corresponding to Figures 3a and b, the entire *x*(*s, t*) curves for the full and quench models. The colors in these panels range from yellow to blue as *t* increases from 2 to 4 in steps of 0.1. Panels (f) and (g) show the same curves as a function of *t*, rather than *s*.

From the fits to the experimental data of *Vorticella* and *Spirostomum* contracting in water (see *SI Appendix Supplementary results* Section A 2), neither organism is expected to obey the quench-like dynamics, being closer to the slow-wave limit. This implies that the organisms are relaxing appreciably *while* the Ca^2+^ wave traverses their lengths.

### D. Scaling behavior distinguishes three dynamic regimes

An additional significant difference between the different models is the scaling of the maximum contraction rate with the viscosity. Notably, this has been studied in previous experimental and computational works on *Vorticella* and *Spirostomum*, against which we may validate our model^9,41,48^.

To explore this scaling behavior, we computed the maximum contraction rate 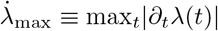 as we varied the non-dimensional parameters *η*_*w,m*_ and *η*_*w,d*_ in the full, quench, and slow wave models. The results are displayed in Figure 3c and d, again setting *w* = 0.1 and *g*_min_ = 0 for simplicity. The dynamics of the quench model only depend on the dimensionless parameter *η*_*m,d*_ (cf. Eq. (15)), and the maximum rate (in units of 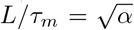) asymptotically scales like 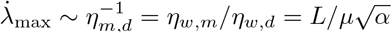. This can be qualitatively understood as follows: as shown in *SI Appendix Supplementary methods* Section B 4, an exact solution for the quench model can be constructed by decomposing the initial condition into spatial Fourier modes and solving for the decay of each mode. Each mode acts like a mass, attached to a wall by a stretched spring with an effective drag constant *η*_*m,d*_, which is suddenly released (see *SI Appendix* Eq. (B64)). By Stokes’ law, the maximum rate should then scale inversely with *η*_*m,d*_ as observed.

For both the full and slow wave models, 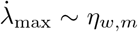 up to a point at which 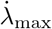 plateaus at 1. This results from the contraction speed not significantly exceeding the speed of the traveling active stretch wave, which is defined as 1 in these units. Interestingly, for both models 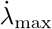 (in units of *L/τ*_*w*_ = *V*) asymptotically scales like 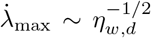. To understand this, recall that the slow wave model can be formally mapped to the diffusion equation with a source term (cf. Eq. (10) and Eq. (13)). The dynamics are thus characterized by the “diffusion constant 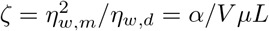. For diffusion, it is known that an initial perturbation to the system spreads out over time with a length scale that grows like 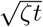, which implies that the corresponding rate scales like 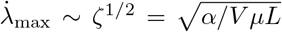. This fact explains both the observed scaling 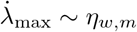 up to its saturation at 1, and the asymptotic scaling 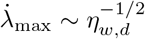. We emphasize that in Figure 3, we have assumed the wave width *W* = 0.1 *L* scales with the length *L*; as demonstrated in *SI Appendix* Figure 8, different scaling behaviors result if one varies the uncontracted length *L* while holding the wave width *W* fixed. In *SI Appendix* Figure 10 we show that this predicted scaling fits our experimental data for *Spirostomum*.

In Refs. 41 and 48, the scaling of 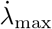 with the solution viscosity *ν* for *Vorticella* is measured to be 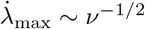. This is consistent with our observed scaling 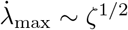 for the full and slow wave models when holding all other parameters constant and only varying *μ* (which is proportional to *ν*). Our model thus exhibits the scaling observed in existing experimental data for *Vorticella*. We note that an alternative description for the *ν*^*−*1*/*2^ scaling based on power-limited contraction was provided in Ref. 41. Our explanation here does not involve a physical argument about power conservation and follows solely from the dynamical equations. It may be possible to show that the energy-based argument is equivalent to the dynamical one presented here, but doing so would likely require a more detailed treatment of thermodynamics of the Ca^2+^ wave (as in Ref. 66), and we leave this to future work.

Scaling data for *Spirostomum* are available in Refs. 8 and 9 (Figure 5b). In these experiments, to aid imaging, the cells are confined in either a narrow agar channel (blue points) or between a microscope slide and a glass coverslip (orange points). Depending on the confinement method, the scaling appears consistent with either 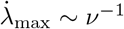, corresponding to the quench regime, or 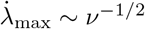. Our fits in Figure 2, together with observation of the asymmetric contraction dynamics (Figure 5a), suggest that *Spirostomum* should not be in the quench regime. The quench-like scaling may be an artifact of using pressure on the coverslips to prevent the organisms from swimming out of view; we speculate that this pressure could hinder contraction during the traversal of the wave, effectively quenching the springs. The imaging rate in Refs. 8 and 9 (800 Hz) may also be too low to resolve the maximum in the contraction rate profile precisely. We thus performed our measurements of the contraction of free-swimming *Spirostomum* using a high-speed camera, recording at a minimum of 2,000 Hz. The results, displayed in Figure 5c, are consistent with the predicted scaling 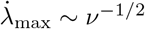, though we cannot conclusively exclude the scaling 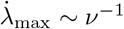. We note that the low 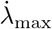 at 1 cP is consistent with the predicted roll-off at low viscosity shown in Figure 3d.

**FIG. 5.**
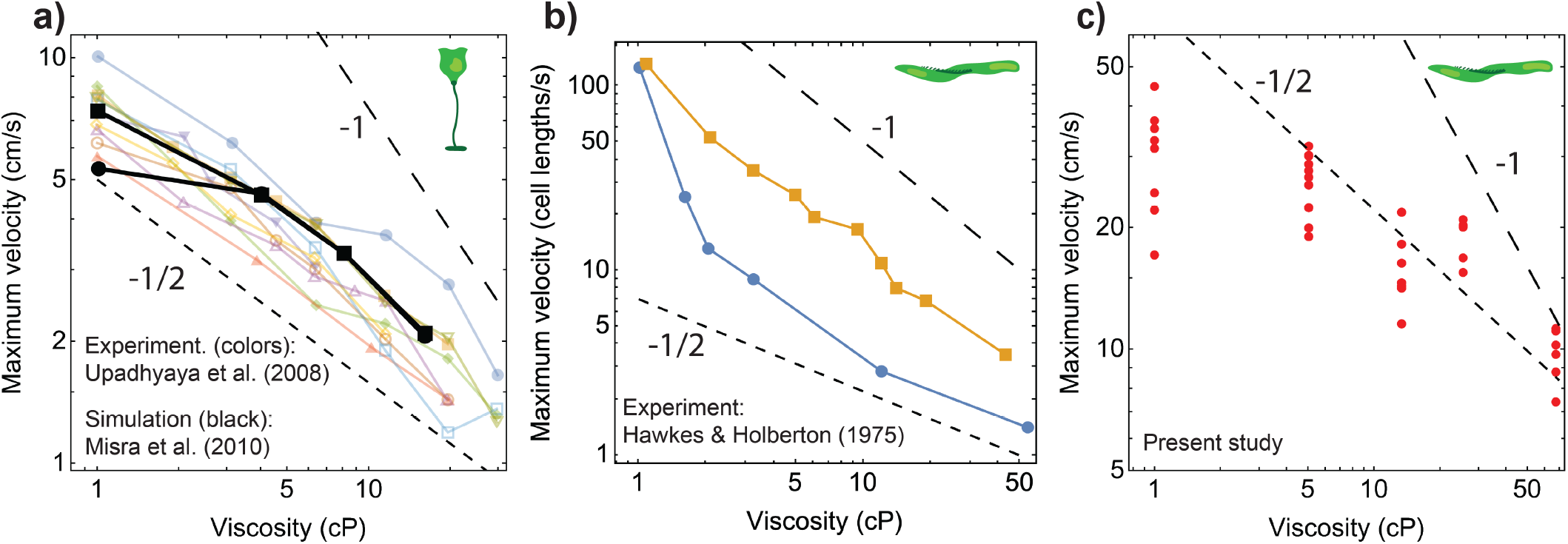
Experimentally measured contraction rate scaling. (a) Published data on the maximum contraction rate of *Vorticella* in solutions of different viscosities. Each colored line represents a single cell exposed to varying viscosities; these data are from Figure 3b of Ref. 41. The black points represent results from simulations, reproduced from Figure 4a in Ref. 48. The circle and square represent two values of the elastic stalk’s Young’s modulus. (b) Published data from Figure 2 of Ref. 9 on the maximum contraction rate of *Spirostomum* in different viscosities and confinement methods: orange represents a setup using plane parallel glass coverslips and blue represents a setup using an agar channel. Each point represents an average of 10-23 measurements, and connecting lines are drawn to guide the eye. (c) Our measurements of the maximum contraction rate of free-swimming *Spirostomum* in solutions of different viscosities. The published data were extracted from Refs. 41, 48, and 9 using WebPlotDigitizer^65^.

The roll-off corresponds to an approach toward a third dynamic regime, which occurs when the active stretch wave is sufficiently slow, or the springs can sufficiently overcome the drag resistance to follow the active stretch wave very closely, with little deviation from equilibrium. In this “quasi-static” limit, there is no dependence of 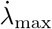 on either the stiffness or drag since the contraction speed is solely determined by the speed of the active stretch wave. This regime occurs in the slow wave model when *ζ* ≫ 1, explaining the approach toward 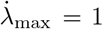 for large *η*_*w,m*_ and small *η*_*w,d*_ in Figures 3c and d. The three scaling regimes—the quench regime 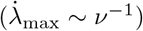, the intermediate regime 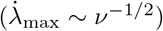, and the quasistatic regime 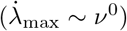—fully characterize the qualitative dynamical behavior of myoneme contraction.

To summarize the dynamical behavior of the system as a function of *η*_*w,d*_ and *η*_*w,m*_, in *SI Appendix Supplementary methods* Section B 5 we introduce the dimensionless quantity *χ*, which measures the peak elastic energy stored in the system as a fraction of the maximum possible energy which is achieved in the quench limit. In Figure 3e we plot *χ* over *η*_*w,m*_∈ [1*/*32, 32] and *η*_*w,d*_ ∈ [1*/*4, 256], and In *SI Appendix* Figure 7 we show corresponding trajectories of the energy and length for these conditions. Let us imagine fixing all parameters except the spring stiffness and the viscosity. Figure 3e then illustrates the system’s ability to respond to the active stretch wave under different mechanical and environmental conditions: for small *η*_*w,m*_, corresponding to soft springs, the system stores a large fraction of the possible amount of internal energy (*χ* ≈ 1), but this fraction decreases as the drag, proportional to *η*_*w,d*_, decreases. By contrast, for large *η*_*w,m*_, the springs closely follow the active stretch wave and never accumulate much internal energy (*χ*≈ 0), and the amount that does accumulate decreases as the drag decreases. This energetic perspective thus provides a unified way to understand the dynamic regimes for organisms using Ca^2+^-based motion.

**FIG. 6.**
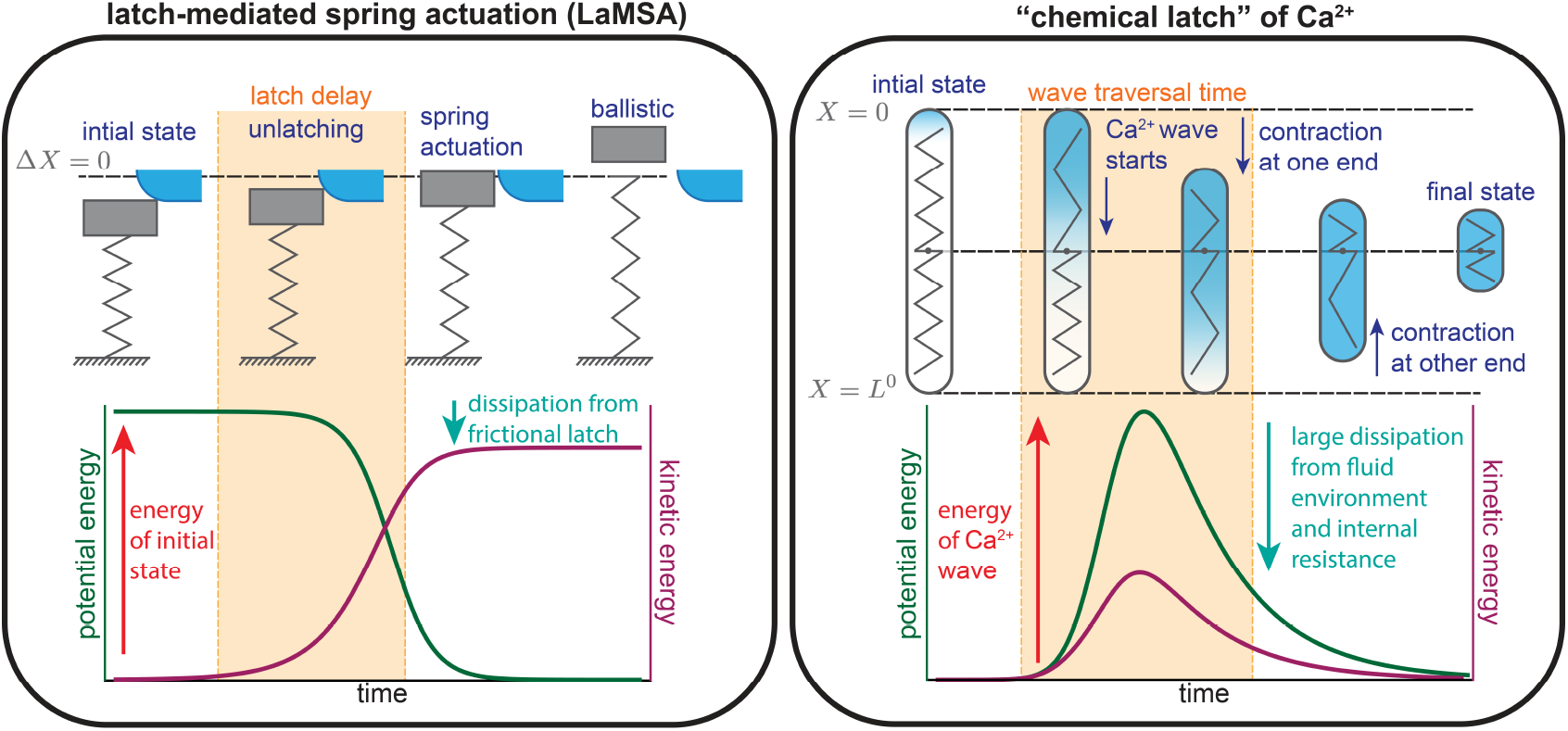
Schematic illustration of a chemical latch. The left panel, adapted from Ref. 67, illustrates a mechanical latch- and-spring system. A latch, shown in blue, slides frictionally off of a compressed spring, shown in gray, which allows for a fast release of the stored potential energy. The finite time during which the latch slides is highlighted in orange. The potential and kinetic energies of the system are schematically plotted at the bottom. At smaller scales, such as the scale of a single-cell organism like *Spirostomum*, a different paradigm accounts for the chemical nature of mechanical activation (via a traveling Ca^2+^ wave). This is illustrated in the right panel, where analogous features of the dynamics in the macroscopic case are shown in the corresponding colors. The traveling wave is shown as a blue gradient, which controls the local rest length (indicated as the number of ridges) of internal springs, shown in gray.

## IV. DISCUSSION

### Summary

We present a minimal mathematical model of myoneme contraction that captures the kinematic features of the organisms *Spirostomum* and *Vorticella*, which exhibit different shapes, sizes, and myoneme arrangements. This model suggests that a fast-traveling wavefront of active stretch, underpinned by rapid Ca^2+^ kinetics, synchronizes contraction across the organism’s length. Using an auxiliary reaction-diffusion model for the Ca^2+^ dynamics, we derive scaling relationships for the speed and width of the active stretch wave that enters the mechanical model. These parameters dictate the rate of chemical driving of the myoneme network. Through non-dimensionalization, we determine the minimal independent set of parameters controlling contraction and further examine two limiting cases: a quench limit and a slow wave limit. The latter approximately applies to both *Spirostomum* and *Vorticella* in water, as confirmed by fitting the model to experimentally obtained length trajectories. We formally map the dynamics in this limit to the well-studied diffusion equation, with the unitless parameter *ζ* playing the role of the diffusion constant. The key geometric and mechanical factors that control contractile dynamics include length, stiffness, wave speed, and drag on the organism. These factors manifest through various non-dimensional combinations in the three dynamical regimes outlined above. Our model analysis reveals three dynamic regimes in which myoneme contraction can occur, each with distinct kinematic signatures, such as the symmetry of contraction about the organism’s midpoint and the exponent with which the maximum contraction rate scales with viscosity. We further differentiate these regimes by a single quantity representing the peak elastic energy stored in the organism relative to the maximal possible value attained in the quench limit.

### Biological implications of dynamic regimes

It is worth-while to consider the possible biological implications of the three dynamic regimes. In the quasi-static regime, the myoneme contractile apparatus remains at mechanical equilibrium while the Ca^2+^ wave traverses the organism’s length. In this limit, the contractile speed is fully determined by the wave speed *V* and does not scale with other parameters. The dissipation of chemical energy is minimized at the expense of the speed of motion. The marine protist *Acantharia* reportedly has myoneme-based spicules that are used to regulate buoyancy by contracting in a Ca^2+^-powered process that is two orders of magnitude slower than contraction in *Spirostomum* or *Vorticella*^18,28^. In this case, contraction may occur quasi-statically. Conversely, the quench regime attains the highest contractile velocities. These high speeds consequently dissipate the most energy because the myoneme apparatus deviates the furthest from mechanical equilibrium. This regime would be suited to processes in which large speeds are physiologically important. For example, *Spirostomum* excites contraction of nearby organisms through “hydrodynamic trigger waves,” requiring it to move sufficiently rapidly to disturb the fluid, dissipating energy in the process^10^. The intermediate regime, where we place the contraction of *Vorticella* and *Spirostomum*, is expected to achieve a balance between thermodynamic efficiency and biological functionality. While myoneme-based systems are predominantly found in ciliated protists, similar ATP-independent contractile protein networks exist in a wide range of other organisms across taxa, facilitating a diverse set of biological functions^8–10,14–26^. If similar mechanochemical dynamics apply in those organisms, then various selection pressures may favor a particular regime for each function.

### Comparison to traditional latch-mediated systems

The theory of latch-mediated spring actuation (LaMSA) provides a general description of motion characterized by a fast unlatching event that releases stored elastic energy^67,68^. It has been applied to analyze macroscopic biomechanical processes like the striking of a Mantis shrimp’s claw and a human finger snap^69,70^. One may expect that LaMSA also explains myoneme-based contraction, but here energy is stored in the form of a difference in Ca^2+^ chemical potential between the cytosol and the endoplasmic reticulum, and release of a “chemical latch” corresponds to initiation of a traveling wave of Ca^2+^ ions. In addition to this difference in the energy storage medium, LaMSA is designed to apply to macroscopic systems, neglecting the strong dissipative forces which are unavoidable at the cellular level. Furthermore, in myonemes, contraction occurs over an approximately continuous spatial domain of many small elastic elements connected together, compared to the discrete mechanical elements considered in LaMSA. These qualitative features of traditional LaMSA systems compared to the myoneme-based “chemical latch” are illustrated in Figure 6. Outside the quench limit, the Ca^2+^ wave, which loads mechanical energy into the system, has a finite traversal time. This means that the system can begin relaxing before the energy has been fully loaded. This is another key distinction from traditional LaMSA systems, which are assumed to be fully loaded before relaxing. LaMSA systems also include a constraint force which is lacking in free-swimming organisms like *Spirostomum*. Exploring the implications of adding constraint forces to myoneme-based systems would be an interesting direction for future research. For example, fixing the ends of *Spirostomum* and then releasing them after the Ca^2+^ wave has passed would allow artificially pushing the contraction process into the quench regime. Although how *physical* latches mediate energy flow has only recently been appreciated in biology^68^, our model offers a guiding framework for how a *chemical* latch dynamically controls and tunes force output in ultrafast single-cell organisms.

### Implications for synthetic biology

Finally, our model offers valuable insights for engineering myoneme-based synthetic systems. Researchers have recently demonstrated the feasibility of reconstituting myonemes *in vitro* and exogenously controlling them using light-patterned Ca^2+^ release^71^. Myoneme-based systems, therefore, provide a mechanism for exerting mechanical forces at subcellular scales, with orthogonal chemistry to light-controllable actomyosin and kinesin-microtubule systems^72–76^. Our model uncovers several critical parameters that can modulate the dynamics of myoneme contraction. For example, increasing cross-linking would enhance the effective *α*, elevating the solution viscosity would raise the effective *μ*, and increasing the effective light-activated Ca^2+^ release rate or reducing Ca^2+^ buffering (thus increasing *D*_*C*_) would adjust *V* and *W* of the active stretch wave. By controlling these experimental parameters, researchers can explore the various regimes identified in this work, paving the way for designing synthetic cells equipped with myoneme-based artificial cytoskeletons that harness the full potential of ultrafast contraction dynamics.

## V. MATERIALS AND METHODS

### A. Ca^2+^ dynamics

The dependence of the active stretch *g*(*S, T*) on the local bound Ca^2+^ concentration may be modeled using the instantaneous linear relation^48^

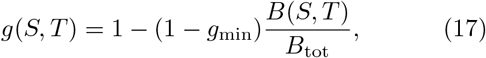

where *B*(*S, T*) is the local concentration of Ca^2+^ bound to myoneme fibers and *B*_tot_ is the total concentration of binding sites (which is assumed to be uniform over the spatial domain). Finding *g*(*S, T*) then requires specifying the dynamics for the bound Ca^2+^ profile *B*(*S, T*). To avoid introducing extra variables, we first analyze the complete dynamics for *B*(*S, T*) by extending a tractable model for a traveling Ca^2+^ wave developed by Kupferman *et al*. in Ref. 57, and we then use results of this analysis to determine an approximate form for *g*(*S, T*) to use in the mechanical model.

The Kupferman model for a traveling Ca^2+^ wave is

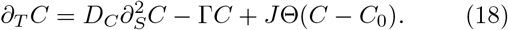

Here, *C*(*S, T*) is the concentration profile of cytosolic Ca^2+^ which diffuses with a diffusion constant *D*_*C*_, leaks from the cytosolic compartment with a first-order rate constant Γ, and is released into the cytosol by internal stores at a spatially dependent rate *J* Θ(*C*− *C*_0_). This last term involves a Heaviside function Θ(*C*− *C*_0_), which approximates the kinetics of calcium-induced calcium release^62^. It “turns on” when the local Ca^2+^ concentration *C*(*S, T*) exceeds a spatially uniform threshold *C*_0_, at which point Ca^2+^ is released into the cytosol at a rate *J*. The authors of Ref. 57 show that the speed of the traveling wave is given by Eq. (5).

We extend the Kupferman model to allow cytosolic Ca^2+^ to bind to the myoneme assembly. This introduces the variable *B*(*S, T*) for the local concentration of myoneme-bound Ca^2+^, and the coupled dynamical equations for *C* and *B* are

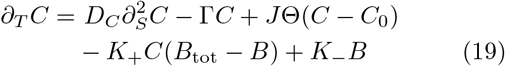

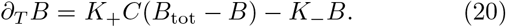

Here, *K*_+_ is the second-order rate constant for binding of cytosolic Ca^2+^ to open myoneme binding sites, which have concentration *B*_tot_ −*B. K*_*−*_ is the first-order rate constant for the subsequent unbinding of myoneme-bound Ca^2+^.

In *SI Appendix Supplementary methods* Section B 2, we analyze Eqs. (B18) and (B19) under mild simplifying assumptions to show that: *i*) the function *B*(*S, T*) is well approximated by a traveling wave with a sigmoidal profile:

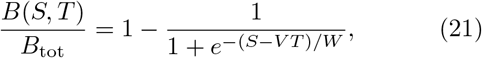

*ii*) its speed *V* is equal to the speed of diffusing Ca^2+^ wave (cf. Eq. (5)), and *iii*) its width *W* can be expressed in terms of the chemical parameters according to Eq. (B31).

Combing Eqs. (17) and (21) gives the following form for the non-autonomous active stretch:

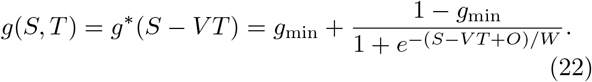

The additional parameter *O* is a length that allows an adjustable offset between *T* = 0 and the time *T* = *O* when the center of the wave arrives at *S* = 0. This allows shifting time to achieve acceptable numerical agreement between boundary and initial conditions, as described in *Materials and methods* Section V C.

### B. Experimental methods

We obtained *Spirostomum* organisms from Carolina Biological and cultured them at room temperature. We prepared culturing medium by mixing one part boiled hay medium solution (Ward’s Science, 470177-390), four parts boiled spring water, and two boiled wheat grains (Carolina, Item #132425) as described previously^35^. We inoculated organisms into fresh media approximately twice per week. For imaging, we pipetted several organisms at a time into microscopy slides with multiple channels (Ibidi, *μ*-Slide VI 0.4) or with multiple wells (Ibidi, *μ*-Slide 8 well). The channels are 400 *μ*m high, allowing the organisms to swim and contract without confinement.

Our *Vorticella* samples were collected from Morse Pond near the Marine Biological Laboratory. We dissected small pieces of plant material to which individual or small groups of *Vorticella* were attached. We added these pieces of plant matter, along with enough imaging medium to cover them, into 8 well coverslip-bottomed sample chambers (Ibidi, *μ*-Slide 8 well).

To alter the viscosity, we prepared stock solutions of Ficoll PM 400 (Sigma F4375) in water and diluted them to concentrations between 3% and 12% (weight per volume). In the case of *Spirostomum*, we pipetted organisms directly into the Ficoll solution. In the case of *Vorticella*, we rinsed the samples several times with the solution within the sample chamber to ensure that the appropriate viscosity was reached. In both cases, slow pipetting was important to prevent shearing of the organisms in the high-viscosity solutions. To calculate viscosity *ν* as a function of percentage *p* of Ficoll in water we used the calibration function *ν* = *e*^0.162*p*^ reported in Refs. 77 and 78.

We performed brightfield imaging on either a Zeiss inverted microscope equipped with a 0.55 NA condenser, a 0.63x defocusing lens, DIC optics, and a 10x air objective or a Nikon Ti2 equipped with a 2x (MRD70020) or 10x (MRD70170) air objective. In both cases, we collected images using a pco.dimax CS camera equipped with at least 2000 and up to 10,000 Hz.

For quantifying *Spirostomum* images, we first used Fiji^79^ to threshold the organisms, and then we used Matlab to run Morphometrics^80^ to obtain the cell contours. We used the pill-mesh option of Morphometrics to obtain cell shape and length. See *SI Appendix* Figure 9 and *SI Appendix Supplementary movies* S3 for a visualization of this mesh generation procedure.

For quantifying *Vorticella* length over time, we manually tracked the position of the end of the stalk attached to the cell body using home-built Python software.

### C. Computational methods

All numerical integration was performed with Mathematica’s NDSolve function, using the method of lines option with a spatial discretization based on tensor product grids. The accuracy and precision goals were set high enough to ensure reasonable numerical convergence. To achieve agreement between the initial conditions Eq. (4) and the boundary conditions Eq. (3), we increased the parameter *O* in Eq. (22) (*o* in the non-dimensional model) so that *g*(*S*, 0) ≈1 across the computational domain. To solve the quench model Eq. (15), we evaluate the exact solution given in *SI Appendix* Eq. (B58) using 50 terms in the Fourier expansion.

## ACKNOWLEDGMENTS

We wish to thank Shane Terrell for help setting up the experiments, Vincent Boudreau for providing samples of *Vorticella* in fresh pond water, Joe Brzostowski, Tanner Fadero, and Sam Lord for technical assistance with the microscopy, KC Huang and Tristan Ursell for providing the Morphometrics software, and Joe Lannan, Christian Pagán-Medina, Joël Lemière, and Jane Maienschein for helpful discussions. This work was mainly supported by National Science Foundation awards EF-1935260, EF-1935261, and EF-1935262. SV was supported by the National Institute of General Medical Sciences of the National Institutes of Health under Award Number R35GM147400. The content is solely the responsibility of the authors and does not necessarily represent the official views of the National Institutes of Health. CF acknowledges support from the University of Chicago through a Chicago Center for Theoretical Chemistry Fellowship. MSB acknowledges funding support from NIH Grant R35GM142588; NSF Grant 1817334; NSF CAREER 1941933; and a gift from the Open Philanthropy Project. FC acknowledges funding support from NIH Grant R35GM141796. We acknowledge the University of Chicago’s Research Computing Center for computing resources, the Marine Biological Laboratory Whitman Summer Investigator Program and the Physiology Course for the use of microscopy equipment, and Zeiss and pco. for loans of cameras.

## Appendix A: Supplementary results

### 1. Supplementary figures

**FIG. 7.**
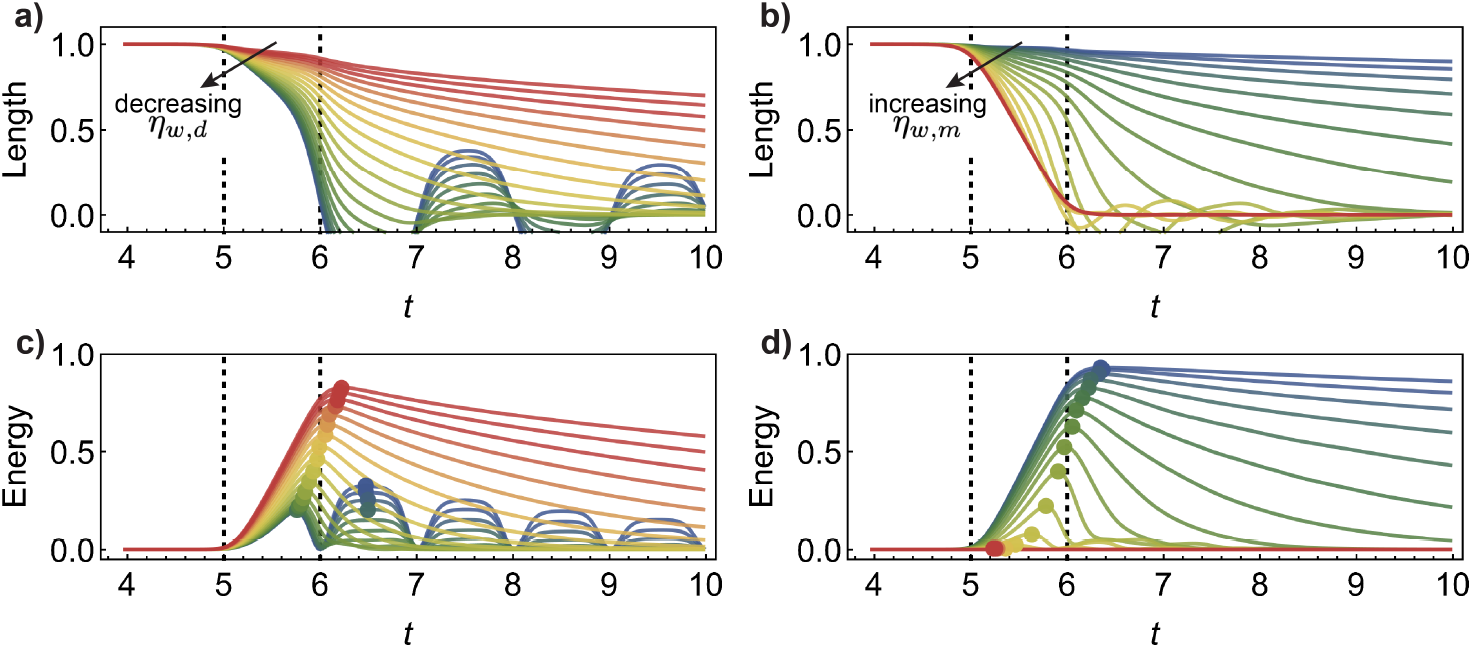
Dependence of length and energy trajectories on key timescales. In panels (a) and (c), *η*_*w,d*_ is varied from 0.25 to 256 in powers of 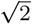 as the colors range from blue to red, with *η*_*w,m*_ = 1 fixed. The dimensionless length *λ*(*t*) and energy *u*_el_(*t*) trajectories are shown; see the main text and *SI Appendix Supplementary methods* Section B 5 for a definition of 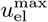. The circles drawn in panel (c) show the values of *u*^max^ which are used to compute *χ* in Figure 3e of the main text (see *SIAppendix* Eq. (B67)). Similarly, in panels (b), and (d), *η*_*w,m*_ is varied from 0.0625 to 32 in powers of 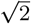 as the colors range from blue to red, with *η*_*w,d*_ = 2 fixed.

**FIG. 8.**
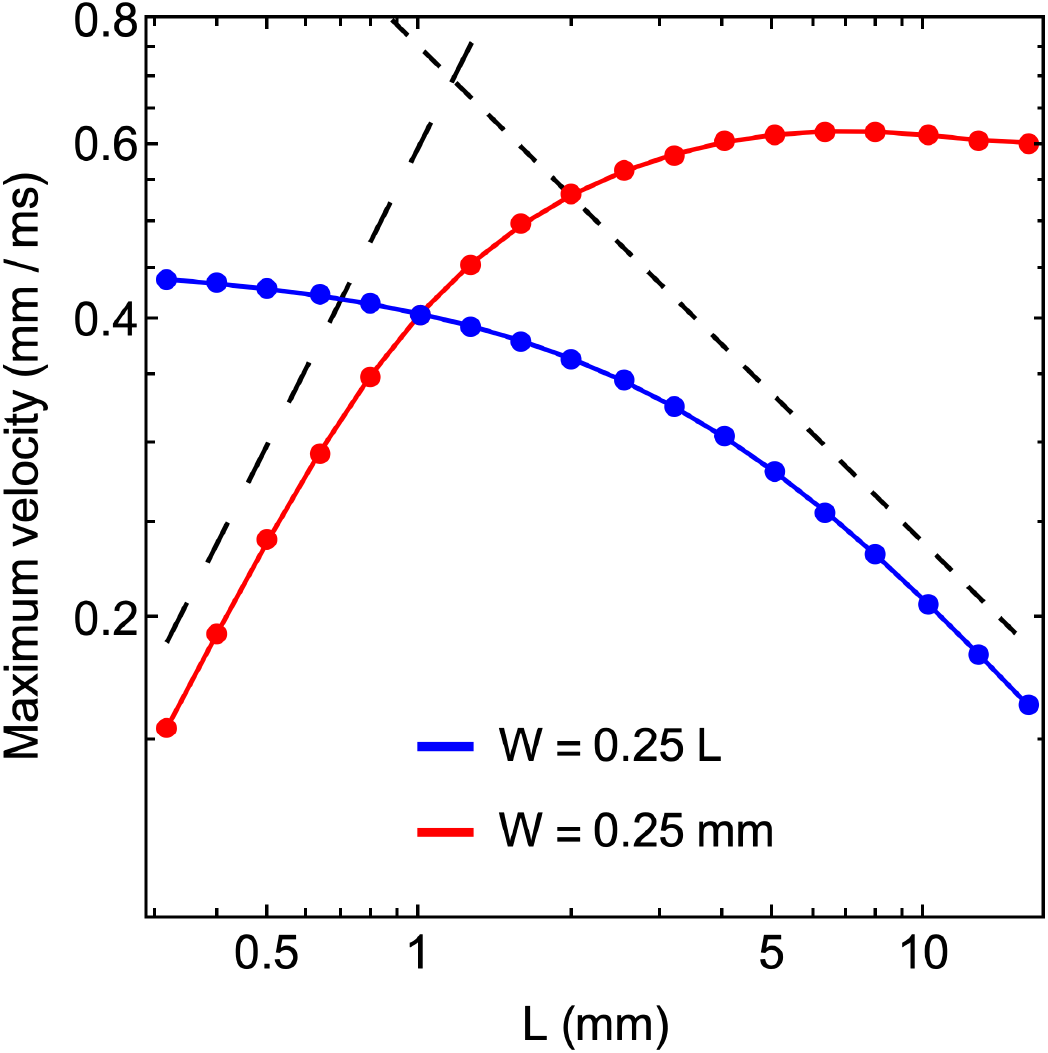
Coupling between length and width affects scaling. In the dimensional model, Eq. (A1) below, the maximum contraction rate 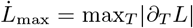 is plotted as we vary the uncontracted length *L*. The red points correspond to conditions in which the width of the active stretch wave is fixed at *W* = 0.25 mm, while the blue points correspond to conditions in which the wave width scales with *L* as *W* = 0.25 *L*. The varying of *W* with *L* qualitatively changes the dependence of 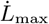. The remaining parameters used for this plot are *α* = 1 (mm*/*ms)^2^, *μ* = 3 ms^*−*1^, *V* = 0.5 mm*/*ms, and *g*_min_ = 0. The short and long dashed lines are proportional to *L*^*−*1*/*2^ and *L*, respectively.

**FIG. 9.**
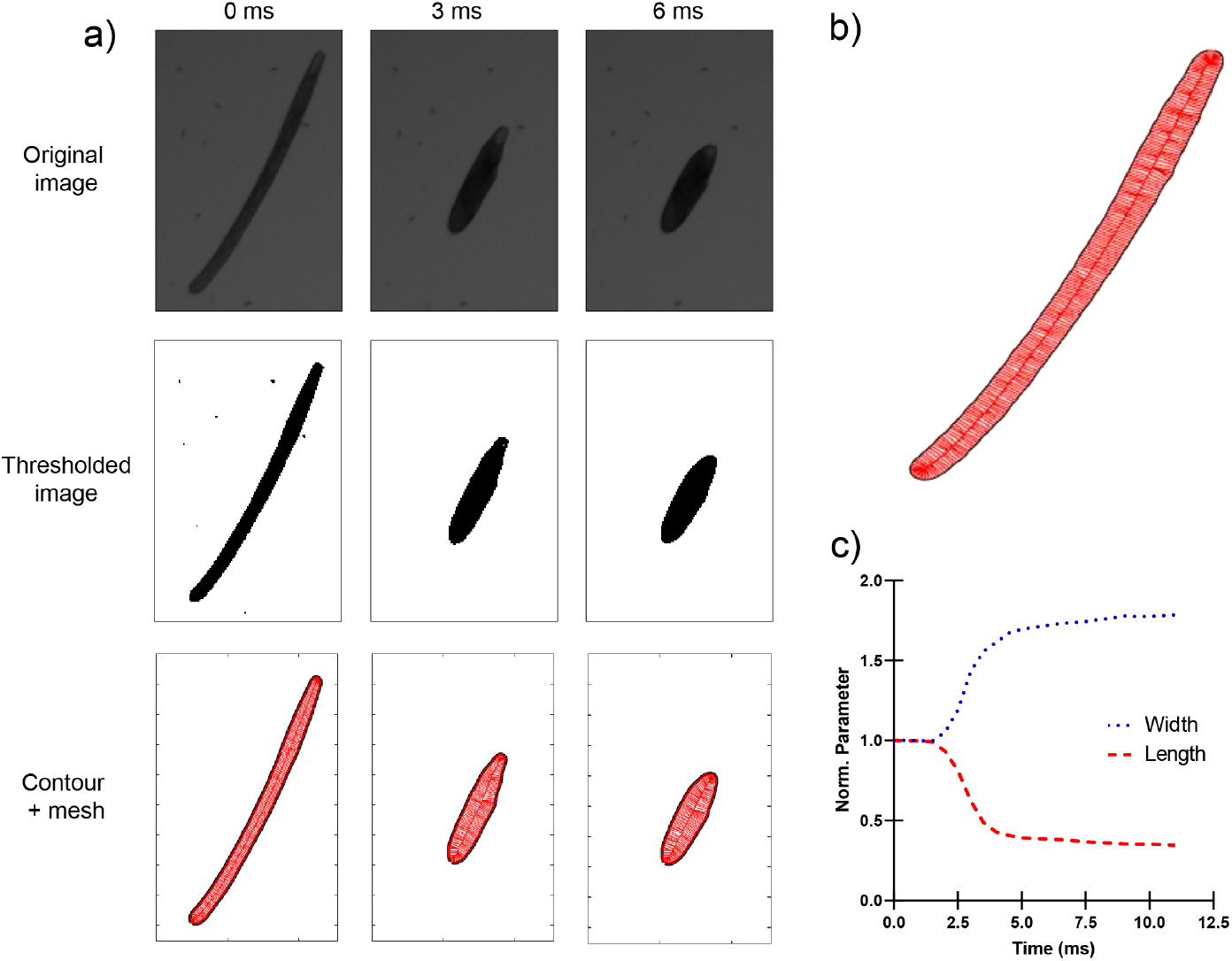
Measuring the length and width of *Spirostomum*. (a) The original video image, the thresholded image, and the detected contours of *Spirostomum* at three moments during contraction. We used FijI^79^ to threshold the images, and we used Matlab to run Morphometrics^80^ with the pill-mesh option to obtain the cell contours. (b) Expanded view of the detected mesh to illustrate greater detail. The instantaneous width is reported as the median value of the width at several points along the organism’s length, and the instantaneous length is reported as that of the central backbone curve. (c) Time course of the length and width, normalized by their initial values.

**FIG. 10.**
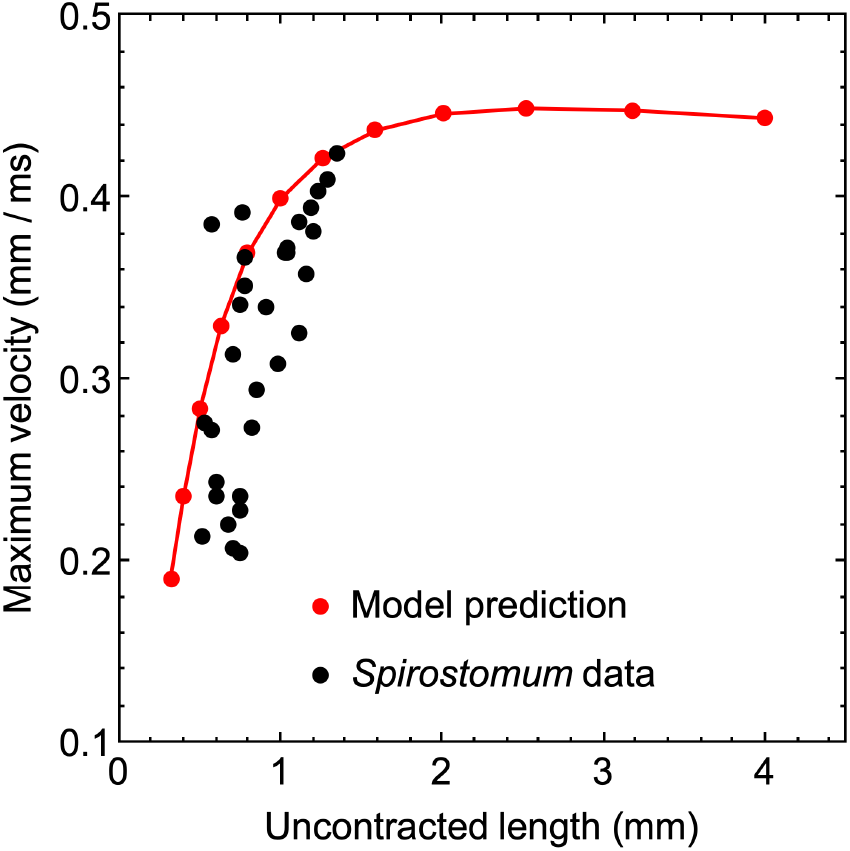
Experimental data for the dependence of contraction speed on initial length of *Spirostomum*. Experimental data on the maximum contraction velocity is plotted against the initial length of the organism. The Pearson correlation coefficient is 0.72 with a p-value of 5 ×10^*−*6^ for this data set. The predicted scaling of maximum contraction velocity with length *L* in the slow-wave limit of the model, Eq. (A2) below, is plotted in red, using the parameters *V* = 0.8 m*/*s, *W* = 0.2 mm, *μ/α* = 5 (mm ms)^*−*1*/*2^, and *g*_min_ = 0.4 (cf. Table I). See Ref. 40 for corresponding data on *Vorticella*.

**FIG. 11.**
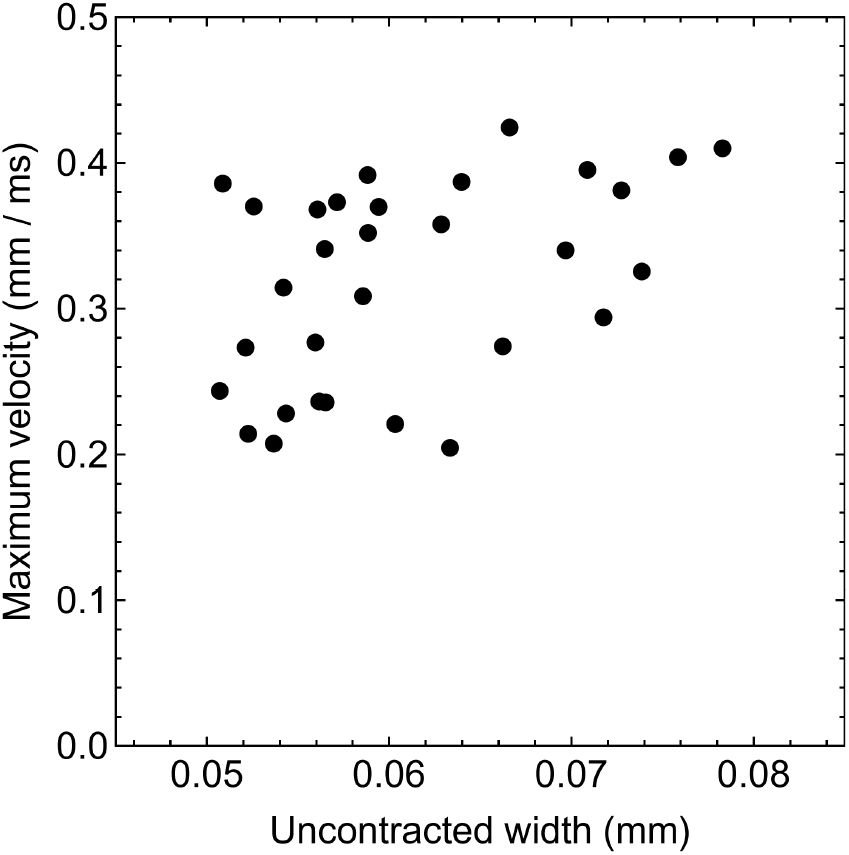
Experimental data for the dependence of contraction speed on initial width of *Spirostomum*. Experimental data on the maximum contraction velocity is plotted against the initial width of the organism. The Pearson correlation coefficient is 0.44 with a p-value of 0.013 for this data set.

### 2. Fitted parameters

Here we fit the model to each of the five length trajectories for *Spirostomum* and *Vorticella* shown in Figure 2a and c of the main text. Figure 12 illustrates the fits to *Spirostomum*, and the fit parameters are given in Table I. The contraction events occur close to the slow-wave limit of the model, in which inertia has nearly dropped out of the dynamics. Consequently, when fitting the full model, we found only the ratio of the parameters *μ* and *α* was constrained, and not their individual values. We thus fit the the curves in two ways. First, we fit the full model including inertia

**TABLE I.** Estimated parameters for *Spirostomum*. The fitted model parameters for the five contraction samples are shown in Figure 12.

| Shared parameters | Sample 1 | Sample 2 | Sample 3 | Sample 4 | Sample 5 |
| --- | --- | --- | --- | --- | --- |
| $L$ (mm) | 0.61 | 0.58 | 0.52 | 0.51 | 0.76 |
| $g_{\min}$ | 0.45 | 0.35 | 0.37 | 0.37 | 0.38 |
| Parameters for full model | Sample 1 | Sample 2 | Sample 3 | Sample 4 | Sample 5 |
| $V$ (mm/ms) | 0.8 | 0.8 | 0.8 | 0.85 | 0.9 |
| $W$ (mm) | 0.11 | 0.12 | 0.1 | 0.13 | 0.13 |
| $\mu/\alpha$ (ms/mm<sup>2</sup>) | 25 | 28 | 45 | 40 | 14 |
| $\zeta = \alpha/V\mu L$ | 0.082 | 0.077 | 0.054 | 0.058 | 0.11 |
| Parameters for slow-wave model | Sample 1 | Sample 2 | Sample 3 | Sample 4 | Sample 5 |
| $V$ (mm/ms) | 0.7 | 0.8 | 0.7 | 0.7 | 0.95 |
| $W$ (mm) | 0.14 | 0.18 | 0.2 | 0.2 | 0.15 |
| $\mu/\alpha$ (ms/mm<sup>2</sup>) | 6 | 5 | 5 | 6 | 5 |
| $\zeta = \alpha/V\mu L$ | 0.390 | 0.431 | 0.549 | 0.467 | 0.277 |

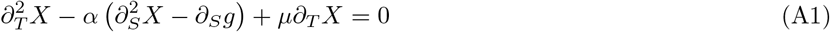

by fixing *α* = 1 and varying *μ*, so that only the ratio *α/μ* is varied independently. Second, we fit the slow-wave model

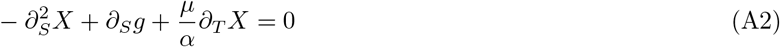

where the parameter *μ/α* appears explicitly. We find that more consistent estimates of the model parameters can be obtained by fitting to the slow-wave model.

We next fit the five length trajectories of *Vorticella* shown in Figure 2c of the main text. Figure 13 illustrates these fits, and the fit parameters are given in Table II.

**TABLE II.** Estimated parameters for *Vorticella*. The fitted model parameters for the five contraction samples are shown in Figure 13.

| Shared parameters | Sample 1 | Sample 2 | Sample 3 | Sample 4 | Sample 5 |
| --- | --- | --- | --- | --- | --- |
| $L$ (mm) | 0.27 | 0.31 | 0.19 | 0.20 | 0.24 |
| $g_{\min}$ | 0.13 | 0.32 | 0.36 | 0.36 | 0.18 |
| Parameters for full model | Sample 1 | Sample 2 | Sample 3 | Sample 4 | Sample 5 |
| $V$ (mm/ms) | 0.3 | 0.23 | 0.13 | 0.13 | 0.13 |
| $W$ (mm) | 0.08 | 0.06 | 0.05 | 0.05 | 0.04 |
| $\mu/\alpha$ (ms/mm<sup>2</sup>) | 30 | 15 | 45 | 45 | 40 |
| $\zeta = \alpha/V\mu L$ | 0.41 | 1.29 | 0.90 | 0.85 | 0.79 |
| Parameters for slow-wave model | Sample 1 | Sample 2 | Sample 3 | Sample 4 | Sample 5 |
| $V$ (mm/ms) | 0.15 | 0.2 | 0.1 | 0.08 | 0.08 |
| $W$ (mm) | 0.06 | 0.08 | 0.05 | 0.04 | 0.04 |
| $\mu/\alpha$ (ms/mm<sup>2</sup>) | 2 | 1 | 1 | 3 | 4 |
| $\zeta = \alpha/V\mu L$ | 12.34 | 16.13 | 52.63 | 20.83 | 12.75 |

To validate these estimates of the parameters, we compare to the parameters reported in Ref. 48 which were used in a detailed computational model of *Vorticella* that achieved good agreement with experiments. We can compare our estimated value for 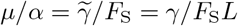 obtained from fitting to the slow-wave model to the corresponding quantity in Ref. 48, where *F*_S_ is the maximal contractile force generated by the myoneme fiber and 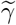 is the drag constant of the organism 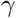 divided by its uncontracted length *L*. In their work, they use the estimate of *F*_S_ = 400 nN measured in Ref. 81. To estimate the drag constant, we assume that drag is dominated by the contribution from the head (with diameter *D*_h_), which is subject to Stokes’ law *γ* = 3*πD*_h_*ν*_w_ where *ν*_w_ = 10^*−*3^ Pa s is the viscosity of water. Using their estimate *L* = 0.2 mm, we obtain the value of *μ/α* reported in Table III. This comparison indicates reasonable agreement between our simplified description of the mechanics and a more detailed model.

**TABLE III.** Estimated parameters for *Vorticella*. The values reported for this study represent the mean and standard deviation of the five samples in Table II.

| Parameter | Present study | Ref. 48 |
| --- | --- | --- |
| $L$ (mm) | $0.25 \pm 0.05$ | 0.2 |
| $g_{\min}$ | $0.27 \pm 0.11$ | 0.25 |
| $V$ (mm/ms) | $0.12 \pm 0.05$ | 0.1 |
| $\mu/\alpha$ (ms/mm<sup>2</sup>) | $2.2 \pm 1.3$ | 5.9 |

We note that the importance of inertia can be estimated by comparing the maximal contraction rate (∼0.1 m*/*s for *Vorticella*) to the expected speed of sound *c*_s_ in the myoneme. As a lower bound for *α* = *F*_S_*/Aρ*, we can estimate the cross-sectional area *A* of the myoneme fiber as that of the whole stalk (∼20 *μ*m^2^) and we take the density of the material as that of water, *ρ* = 1 kg*/*m^3^. With this, we find *α* ∼20 m^2^*/*s^2^ and *c*_s_ ≈4.5 m*/*s. A rough estimate of the speed of sound is thus at least an order of magnitude faster than the maximal contraction speed. This further suggests that for the conditions in which we measured contraction the organisms are in the slow-wave limit in which inertia can be neglected.

## Appendix B: Supplementary methods

### 1. Derivation of continuum model

Here we derive the continuum model of myoneme contraction, Eq. (2) in the main text. We treat the system as a one-dimensional chain of springs with open boundary conditions (or one end fixed for *Vorticella*) and non-autonomous rest lengths for each spring. We consider a set of *N* masses connected by *N*− 1 springs, where the force on the *i*^th^ mass is given by

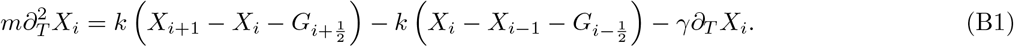

Here, *m* is the mass, *X*_*i*_(*T*) is the horizontal position of mass *i, T* is time, *k* is the stiffness of every spring, 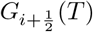 is the current rest length of the spring connecting masses *i* and *i* + 1, and *γ* is the drag coefficient. We use capital letters for dimensional variables and lowercase letters for dimensionless variables. To pass to a continuum description, we label the material coordinate *S* and let the positions *X*_*i*_ and *X*_*i*+1_ be separated by a material distance *δ* which is assumed equal to the initial rest length 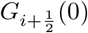 for each *i*. The original material length is *L*, so that *δ* = *L/*(*N* − 1).

We can rewrite Eq. (B1) as

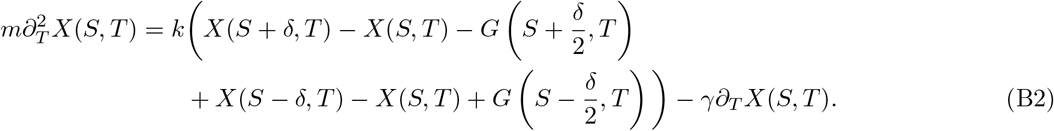

We next assume *δ* is small and expand around *X*(*S, T*) and *G*(*S, T*). To second order in *δ*, this gives (suppressing the dependence on *S* and *T* for clarity)

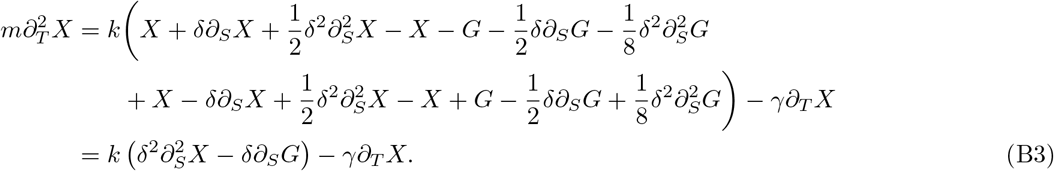

We switch to the dimensionless quantity *g*(*S, T*) = *G*(*S, T*)*/δ* representing the instantaneous rest length *G* divided by the material distance *δ*, which we call the active stretch. This gives

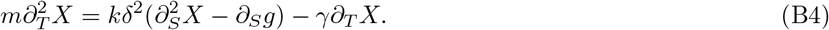

Next we introduce the constants 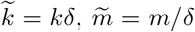, and 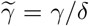, which represent how the stiffness, mass, and drag coefficient physically scale with the discretization length *δ*, and then take the limit *δ* →0 such that all third and higher order terms in the expansion vanish. This gives

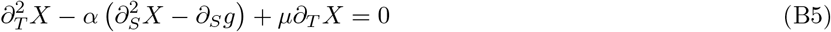

where 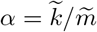 and 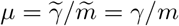. The parameter *α* has units of velocity squared, and the parameter *μ* has units of inverse time.

We assume that at *T* = 0 the length of the organism linearly increases up to its undeformed material length, such that

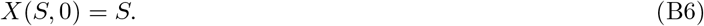

We also let it be at rest initially, such that

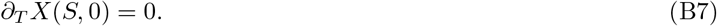

We next consider the boundary conditions for Eq. (B5) at the edges of the domain, *S* = 0, *L*. In the case of *Spirostomum*, we take stress-free boundary conditions at both ends, while for *Vorticella* we use the stress-free condition at *S* = 0 and the Dirichlet condition *X*(*L, T*) = *L*. To derive the stress-free boundary condition, we repeat the discretization steps given above at the mass *i* = 0. We start with

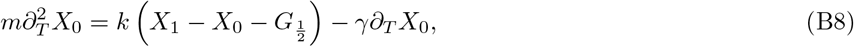

and, passing to the continuous material coordinate *S*,

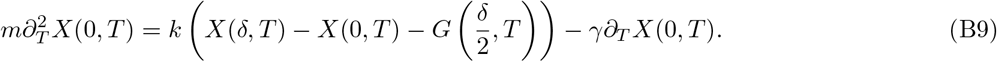

Upon expanding with respect to *δ*, we find (again dropping explicit argument dependence)

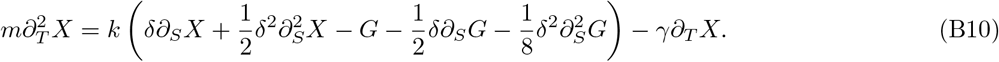

Introducing again *α, μ*, and *g*, we have

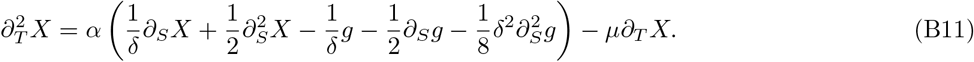

Taking the limit *δ* → 0 leaves us with the following condition at *S* = 0:

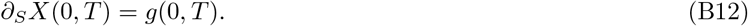

Note that the drag and inertial contributions to the force balance have not survived the limit in this case because the terms proportional to *∂*_*S*_*X* and *g* do not cancel out in the expansion as they did in Eq. (B3), and these terms dominate as *δ* →0. Through the same steps (or by symmetry considerations), we obtain an identical stress-free condition applying at *S* = *L*.

### 2. Description of Ca^2+^ model

#### a. Review of Ca^2+^ models

There is a wide range of mathematical models that describe the time and space-dependent dynamics of intracellular Ca^2+^, reviewed in Refs. 61 and 62. These dynamics are quite varied, including concentration patterns that are homogeneous in space but oscillatory in time, outward-propagating spirals, and the fast-traveling wavefronts which underlie mechanical contraction in *Spirostomum* and *Vorticella*^29,48–51^. An important process giving rise to these dynamics is calcium-induced calcium release (CICR), in which binding of Ca^2+^ to IP_3_ and ryanodine receptor calcium channels causes them to release additional Ca^2+^. This positive feedback does not continue indefinitely, as the probability of the channel being open (and hence its release rate) has a bell-shaped dependence on the logarithm of bound Ca^2+61^. This dual role of Ca^2+^ inducing and subsequently inhibiting the channel release gives rise to the oscillatory patterns observed at long timescales. For computing the speed and width of the traveling wave front in our case, however, it is sufficient to consider only the positive feedback.

Models of Ca^2+^ dynamics differ widely in their complexity, with the number of free parameters ranging from just a handful to more than a hundred^61^. Key features that distinguish Ca^2+^ models include:

- The number of separate species tracked (i.e., just cytosolic Ca^2+^ or also endoplasmic reticulum Ca^2+^ and intermediate signaling molecules like IP_3_)
- Whether the channels are modeled as continuously or discretely distributed (with the latter choice found in the “fire-diffuse-fire” models^82,83^)
- The level of detail for the CICR kinetics of Ca^2+^ channels, which are multimeric molecules that cooperatively bind Ca^2+^ and signaling molecules

With the goal in mind of constructing a minimal model of *Spirostomum* and *Vorticella*, and given the scarcity of experimentally measured kinetic parameters for these systems, we consider a highly simplified model which allows for traveling wave solutions and has clear interpretable parameters. Additionally, we opt for a continuous distribution of Ca^2+^ receptors (rather than a discrete fire-diffuse-fire description), as it is significantly more tractable. In the fire-diffuse-fire models, the continuous case is formally obtained as a limit of the discrete case when the release rate of Ca^2+^ is rate-limiting rather than the inter-receptor diffusion time^82,83^.

#### b. Kuperfman model

For these reasons, we use the Kupferman model of Ref. 57 as the basis for our analysis of Ca^2+^ dynamics and its mechanical effects in *Spirostomum* and *Vorticella*. It tracks the cytosolic Ca^2+^ concentration as a function of time *T* and one space dimension *S*. Below, we also consider the concentration of myoneme-bound Ca^2+^. In the laboratory frame, the reaction-diffusion dynamics for the cytosolic Ca^2+^ concentration *C* are

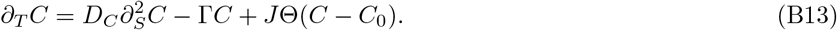

Here, *D*_*C*_ is the Ca^2+^ diffusion constant in the cytosol, Γ is the first-order rate constant for Ca^2+^ leakage out of the cytosol, *J* is the rate of Ca^2+^ release by the open channels, and *C*_0_ is the threshold concentration for the channel to open and begin releasing Ca^2+^. Note that in this model of CICR, all of the complexity of cooperative binding of Ca^2+^ to the channels, dependence on local signaling molecule concentrations, and later inhibition of release is removed by assuming that the release rate dependence on Ca^2+^ concentration is captured by a Heaviside function Θ(*C*− *C*_0_) (rather than some non-monotonic function which would be necessary to describe the dynamics on a longer timescale). Eq. (B13) is rescaled in two steps: first, we non-dimensionalize and we then shift to the frame that is moving together with the traveling wave. The new non-dimensional variables *c, t*, and *s* are introduced by expressing concentration in units of *C*_*u*_ ≡ *C*_0_, time in units of *T*_*u*_ ≡ *C*_0_*/J*, and length in units of 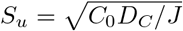. The traveling frame is found by introducing the variable *u* = *s− vt*, where the traveling wave speed *v* is expressed in units of 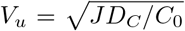 With this rescaling, Eq. (B13) becomes

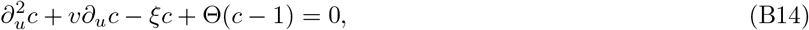

where *ξ* ≡ *C*_0_Γ*/J*.

Eq. (B14) can be solved exactly, which is a key benefit of the Kupferman model^57^. The solution is

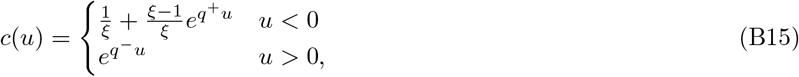

where

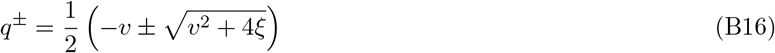

and

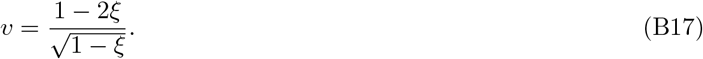

The point *u* = 0 is the boundary between the open (*u >* 0) and closed (*u <* 0) channels, such that at *u* = 0 we have *c* = 1. To obtain this solution to the ODE Eq. (B14) on the domains (−∞, 0) and (0, ∞), the free constants were determined by the matching conditions *c*(0^*−*^) = *c*(0^+^) = 1 and *∂*_*u*_*c*(0^*−*^) = *∂*_*u*_*c*(0^+^).

#### c. Width of myoneme-bound Ca^2+^ profile

We next extend the Kufperman model by introducing the concentration *B*(*S, T*) of Ca^2+^ bound to myoneme proteins. Let *B*_tot_ be the conserved total concentration of myoneme binding sites, which is spatially uniform. When calcium totally saturates the available binding sites on myoneme, then *B* = *B*_tot_. The new reaction-diffusion dynamics for *C* and *B* are

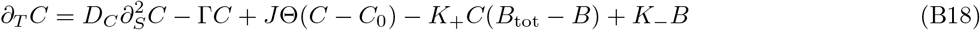

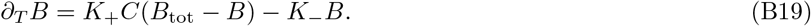

Here *K*_+_ is the dimensional binding rate and *K*_*−*_ is the dimensional unbinding rate. We have assumed that the bound Ca^2+^ does not diffuse. By numerically integrating Eq. (B18) and Eq. (B19), we observed that the traveling profile of *B* has a width that depends on several factors including the binding rate *K*_+_ (see Figure 15 below). The width in turn affects the width of the stretch profile *g*, which can be modeled as a linear function of *B*. The width of the stretch profile *g* is an important feature governing the contraction dynamics in the mechanical model (see Figure 2e in the main text). Our goal here is to study how the width of the *B* profile depends on the kinetic parameters in Eq. (B18) and Eq. (B19).

**FIG. 12.**
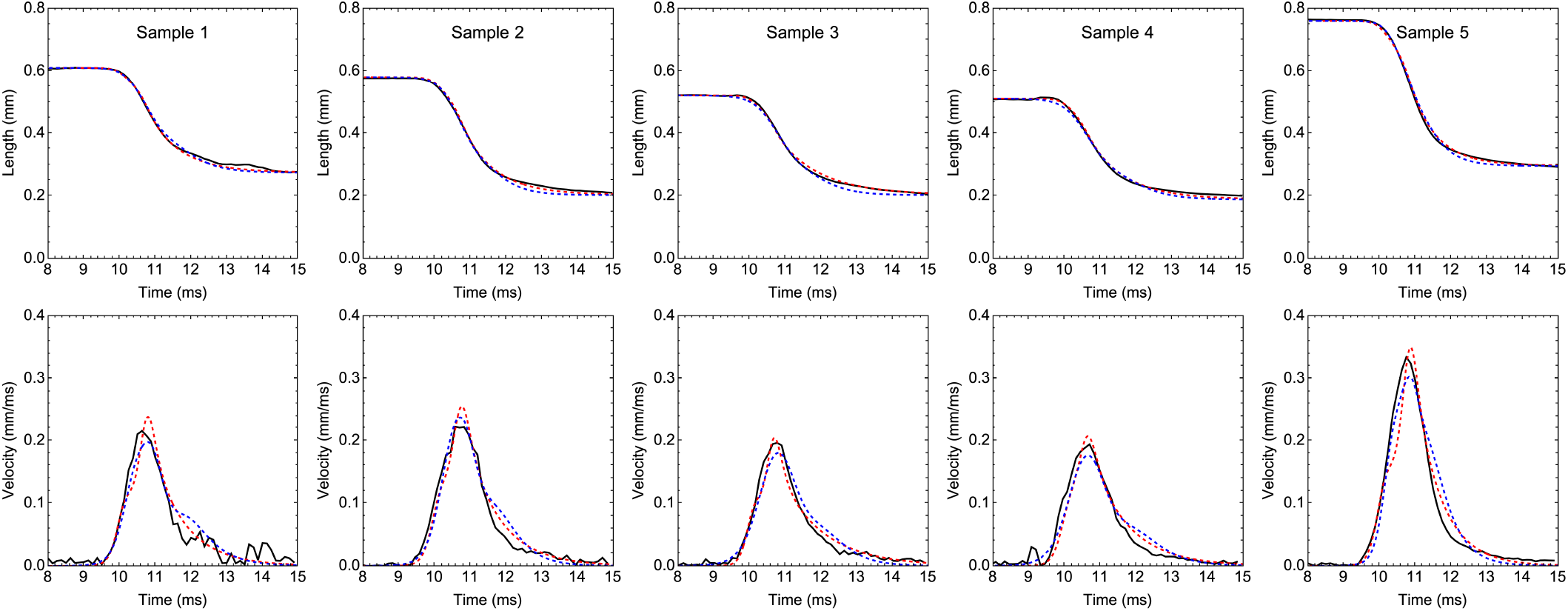
Fits for *Spirostomum*. Experimental trajectories of length and contraction rate are shown as black curves, and fitted model solutions are shown as dashed red (full model) and blue (slow-wave model) curves for the five samples in Figure 2a and b of the main text. Due to experimental limitations, we did not track which traces are from multiple contractions of the same organism versus from different organisms.

The rescaled versions of these equations are

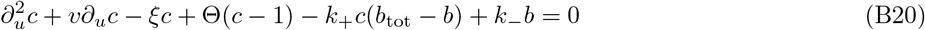

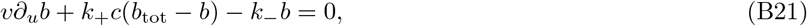

where 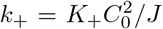, *k*_*−*_ = *K*_*−*_*C*_0_*/J, b* = *B/C*_0_, and *b*_tot_ = *B*_tot_*/C*_0_. Unfortunately, Eq. (B20) and Eq. (B21) are analytically intractable due to the non-linearity from the binding reaction. We therefore look for an approximate solution by relying on two assumptions:

1. The total myoneme concentration *b*_tot_ is sufficiently small that the dynamics of *c* do not depend on *b*.
2. In the range over which *b* is varying, around *u* = 0, the form of *c*(*u*) is approximately linear.

Regarding the first assumption, we find numerically that increasing *b*_tot_ introduces small corrections to the predicted scaling behavior. Nonetheless, we can still gain insight into the model by assuming that the first assumption holds. Regarding the second assumption, we can check graphically that the profile of *c* is indeed roughly linear over the range where *b* is changing (although this also breaks down if *b*_tot_ is large). The linear approximation to *c*(*u*) around *u* = 0 is

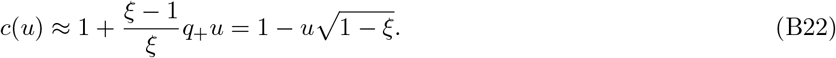

This holds for both sides of the piecewise definition in Eq. (B15) since the derivatives are matched at *u* = 0. In this approximation, *c*(*u*) goes to zero at 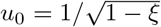, and since *b* cannot be greater than zero if *c* = 0, then *b* must go to zero at that point as well.

With these assumptions the dynamics are reduced to

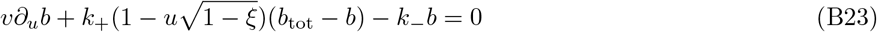

which admits an exact solution:

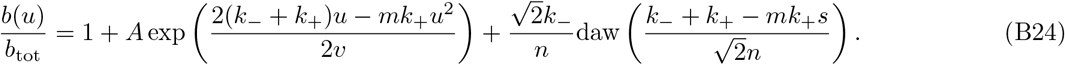

Here, 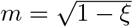 and 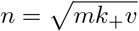, and daw is the Dawson function, defined as

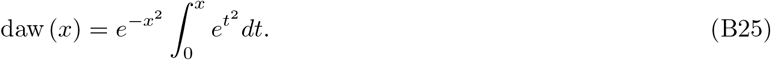

The constant *A* can be fixed by requiring *b*(*u*_0_) = 0, resulting in

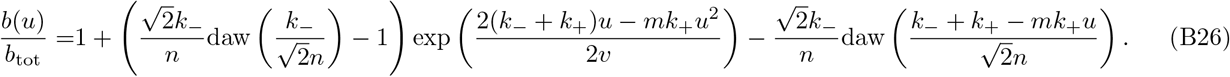

This is defined for *u < u*_0_, while for *u*≥ *u*_0_, *b*(*u*) = 0. Eq. (B26) can be simplified because *k*_*−*_*/n ≪*1 for typical values of the parameters, while the Dawson function does not exceed 1 in absolute value. We can thus write

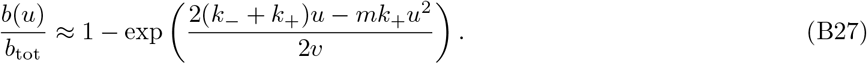

Furthermore, the quadratic part of the exponent dominates after rescaling back to dimensional units, allowing us to finally write

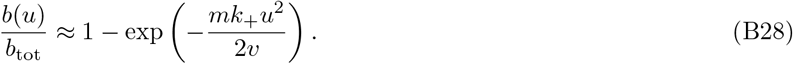

**FIG. 13.**
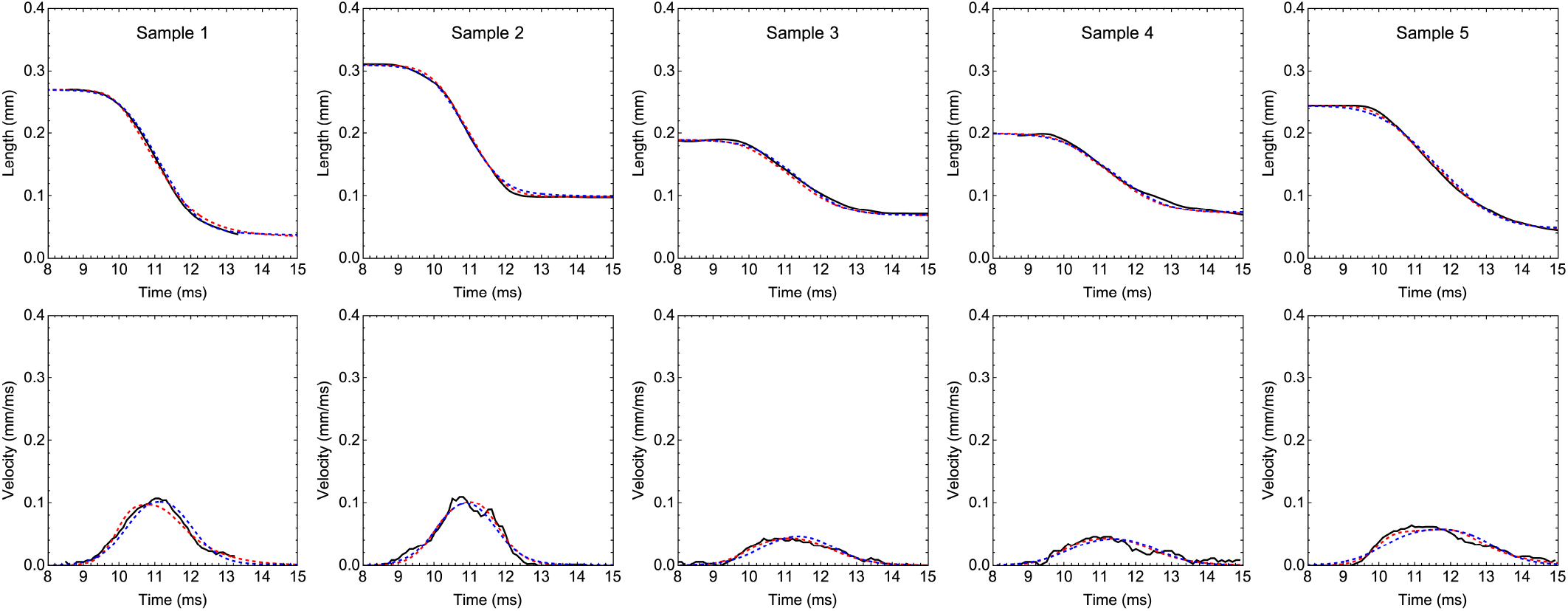
Fits for *Vorticella*. Experimental trajectories of length and contraction rate are shown as black curves, and fitted model solutions are shown as dashed red (full model) and blue (slow-wave model) curves for the five samples in Figure 2c and d of the main text. Due to experimental limitations, we did not track which traces are from multiple contractions of the same organism versus from different organisms.

In Figure 14 we compare Eq. (B26), Eq. (B27), and Eq. (B28) for typical values of the parameters.

**FIG. 14.**
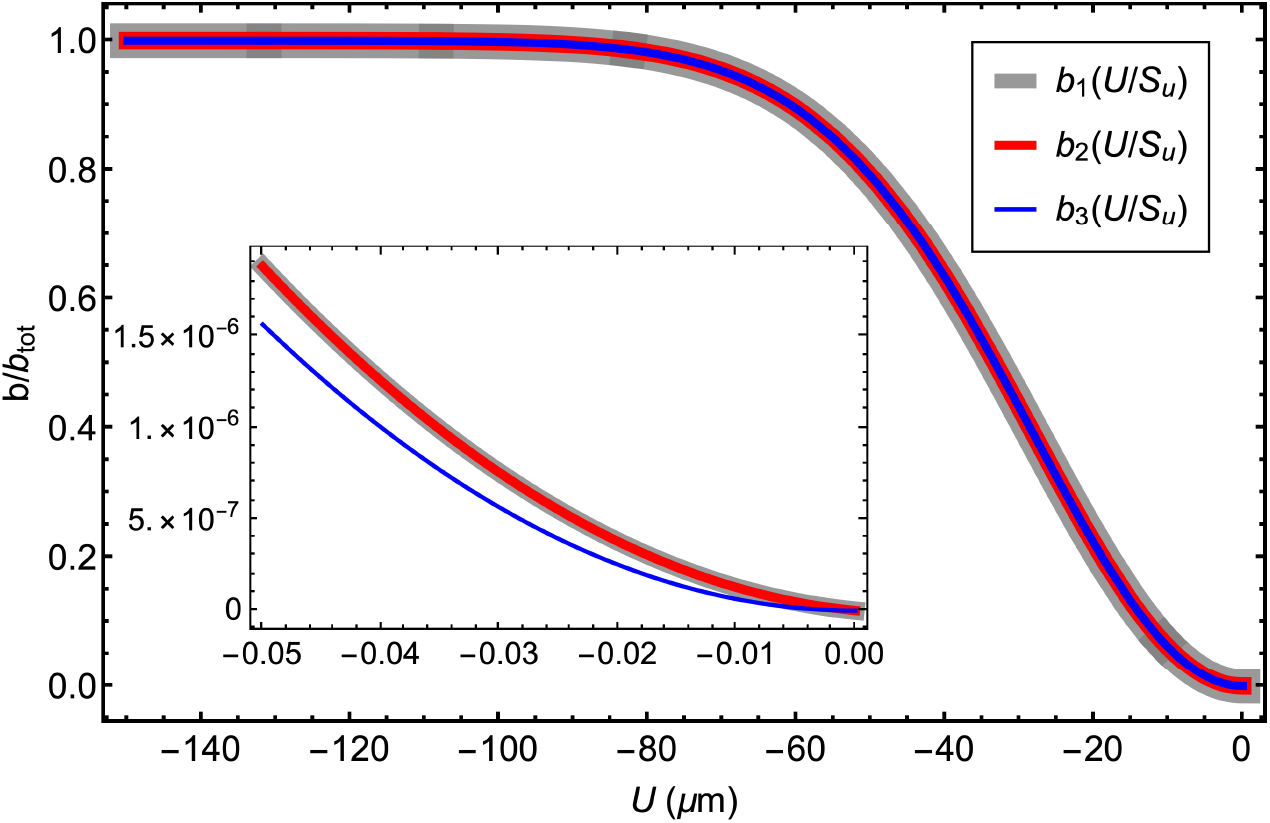
Comparison of Eqs. (B26), (B27), and (B28). For the parameters in Table IV, we show the three equations for *b*(*u*), where *b*_1_(*u*) refers to Eq. (B26), *b*_2_(*u*) refers to Eq. (B27), and *b*_3_(*u*) refers to Eq. (B28). For these parameters, 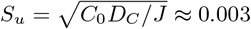. The inset show the region close to the origin, where *b*_3_(*u*) visibly differs from *b*_1_(*u*) and *b*_2_(*u*).

Eq. (B28) is a Gaussian function of *u* with a width given by 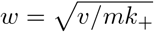. Assuming *m* ≈ 1 since *ξ* is typically small, in dimensional units we have

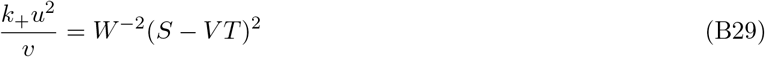

where

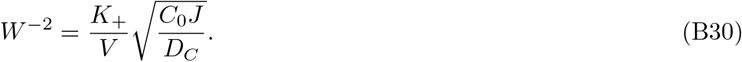

We can simplify using the fact that 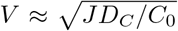 (i.e., *v* ≈1) since *ξ* is small in Eq. (B17) and, by assumption, the dynamics of *c* do not depend on *b*. This gives the final result

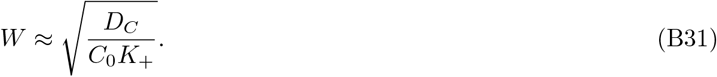

Eq. (B31) predicts that the width *W* only depends on the three quantities *D*_*C*_, *C*_0_, and *K*_+_. We confirm this numerically in the next section.

An interesting extension to the model, which we leave to future work, would be to introduce tension-gated Ca^2+^ channels in the chemical dynamics, to allow for two-way coupling between chemistry and mechanics^10^. This kind of mechanochemical coupling would allow for additional dynamical features like pattern formation, limit cycles, or damped oscillations without requiring inertia. Related work on the mechanochemical dynamics of elastic media can be found, for instance, in Refs. 84 and 85. Another potential extension is to consider a finite delay between Ca^2+^ binding and rest length shrinking, which would introduce an additional timescale to the model. However, we have not addressed this possibility in our current work. Furthermore, we have not explored the possibility that Ca^2+^ binding could alter the local stiffness of the myonemes.

#### d. Numerical results

We demonstrate through numerical tests the accuracy of the above scaling form for *W*, Eq. (B31). Throughout this section and in the inset of Figure 2e in the main text, we use the parameters in Table IV unless noted otherwise. These parameters are mostly in keeping with those used in Ref. 57 except for *K*_+_ and *K*_*−*_, which are based on values roughly inferred for *Vorticella* from Ref. 48.

**TABLE IV.** Kinetic parameters of the Ca^2+^ model.

| Symbol | Value | Units |
| --- | --- | --- |
| $D_C$ | 200 | $\mu\text{m}^2/\text{s}$ |
| $C_0$ | 0.5 | $\mu\text{M}$ |
| $\Gamma$ | 100 | $1/\text{s}$ |
| $J$ | $10^7$ | $\mu\text{M}/\text{s}$ |
| $K_+$ | 0.5 | $1/\mu\text{M s}$ |
| $K_-$ | 0.15 | $1/\text{s}$ |
| $B_{\text{tot}}$ | 1000 | $\mu\text{M}$ |

Using the methods described in *Materials and methods* section V C of the main text, we numerically integrate the full chemical dynamics, Eq. (B18) and Eq. (B19), on a domain of length *L* = 1000 *μ*m for 0.05 s, and we use the boundary condition *∂*_*S*_*C*(0, *T*) = *∂*_*S*_*C*(*L, T*) = 0. We initiate the wave by setting *C*(*S*, 0) = *J/*Γ on the segment *S* ∈[0, 100] and *B*(*S*, 0) = 0 everywhere. To obtain *W* and test the predicted scaling form in Eq. (B31), we fit to the profile *B*(*S*, 0.005 s) the sigmoidal function

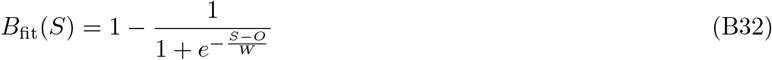

for the offset *O* and width *W*. Note that this functional form does not match the derived form in Eq. (B28) but nonetheless has a well-defined characteristic width. We favor using the sigmoidal approximation to the bound concentration profile because it does not involve a piecewise definition as the form in Eq. (B28) does (where *b*(*u*) = 0 for *u*≥ 0), making it much easier to work with. If the derived scaling is observed for the width of this function, it suggests that the width’s scaling behavior does not depend on the specific functional form in Eq. (B28).

In Figure 15a we show the concentration profiles for *C* and *B* from *T* = 0 s to *T* = 0.025 s, illustrating the propagation of the diffusing and bound Ca^2+^ profiles. We note that the step jump near *S* = 100 on the left of these plots is an artifact of our choice for initial conditions and does not play an important role in the mechanical model of contraction, where a traveling sigmoidal approximation to the bound calcium concentration is used. In Figure 15b we show these profiles at *T* = 0.01 s, along with the fit of Eq. (B32) to *B*(*S*). Note that the profile for *C* is indeed approximately linear over the range where *B* is changing, validating our previous assumption. We show similar profiles for the condition when *K*_+_ = 0.125 (*μ*M *s*)^*−*1^ and *D*_*C*_ = 800 *μ*m^2^*/*s (Figure 15c), and when *B*_tot_ = 16, 000 *μ*M (Figure 15d). In Figure 16 we show the scaling of *W* with the quantities *K*_+_, *K*_*−*_, *D, C*_0_, Γ, and *J*. We predict from Eq. (B31) that the scaling exponents for these will be, respectively, −1*/*2, 0, 1*/*2, −1*/*2, 0, and 0, and these predictions are numerically confirmed.

### 3. Model limitations and extensions

The goal of the the model developed in this paper is to reproduce available experimental observations, such as asymmetrical contraction and the scaling of contraction speed with environmental viscosity, and yet still to be sufficiently simple that we can analytically study its behavior. To this end, we have abstracted several biophysical details into coarse-grained parameters, and we have kept only those features expected to play dominant roles in controlling the contraction speed. These retained features include the elasticity of the myonemes, the Stokes drag on the compression motion, and the dynamics of the chemical wave controlling the myoneme actuation. We have neglected possible volume or area conservation of the cells, spatial variation in the drag coefficient along the cell’s length^41,48^, twisting dynamics or three-dimensional geometry of the cell bodies^35,48^, braking contributed by poroelasticity of the intracellular environment^60,86^ or entanglement of the internal organelles^87^, elastic resistance by auxiliary elastic elements^35^, and intermediate Reynolds number corrections to the viscous drag^41,48^. Detailed treatment of these neglected physical ingredients will likely require organism-specific considerations and numerical approaches, but in the remainder of this section we discuss how some of these features could be approximately included in our modeling framework. We emphasize the limits in which the extended models reduce to our minimal model, which clarifies our assumptions and why our minimal model can successfully capture organismal dynamics.

**FIG. 15.**
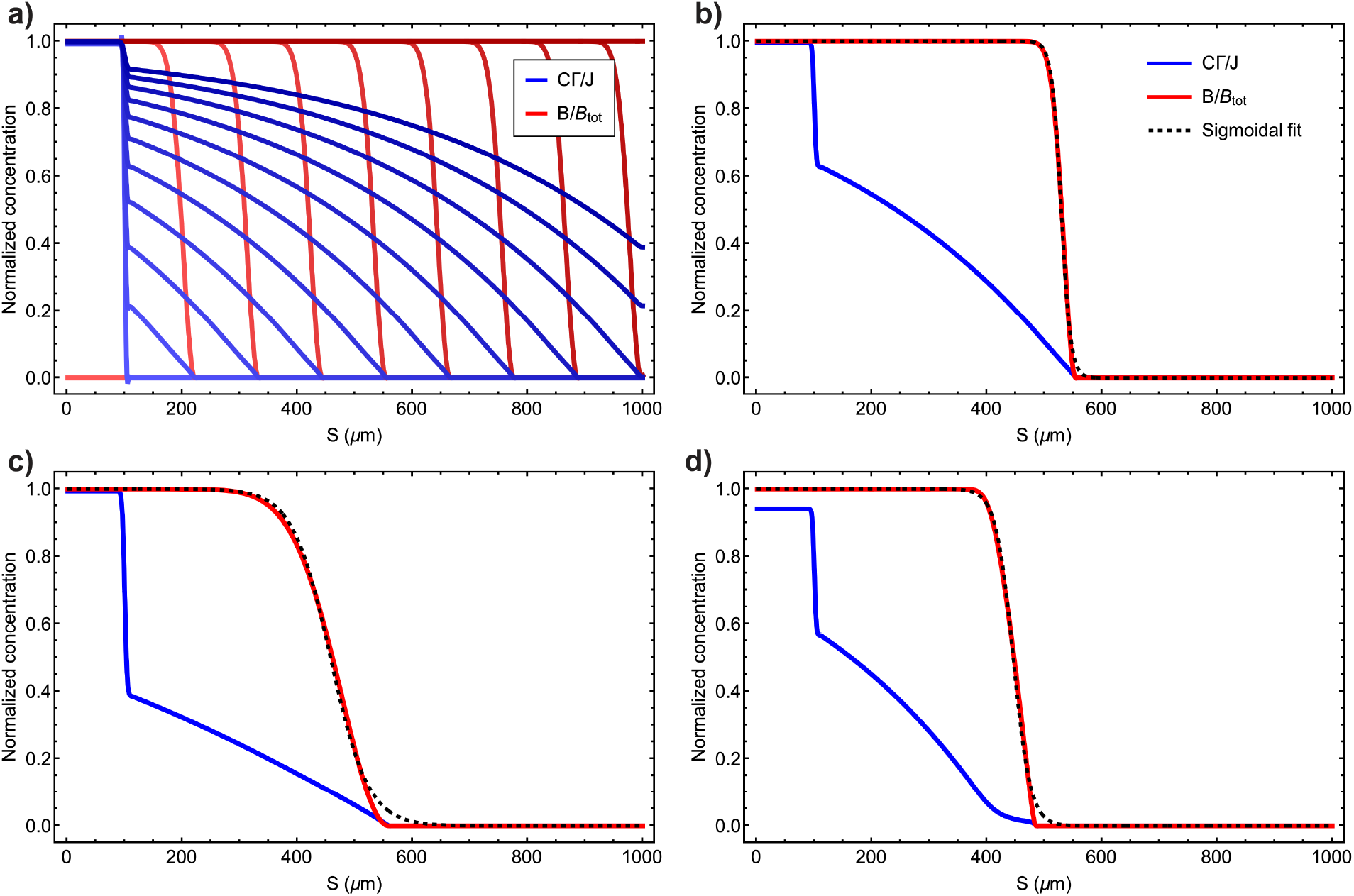
Numerical integration of Eq. (B18) **and** Eq. (B19). (a) The profiles of *C* and *B* from *T* = 0 s to *T* = 0.025 s in steps of 0.005 s as brightness ranges from light to dark. (b) The profiles of *C* and *B* at *T* = 0.01 s. The fit of Eq. (B32) to the profile of *B* is shown as a dashed black line. The fitted width of *B* is 8.2 *μ*m. (c) The profiles of *C* and *B* at *T* = 0.001 s for the same parameters as in (a) but with *K*_+_ = 0.125 (*μ*M*/s*) and *D*_*C*_ = 800 (*μ*m^2^*/s*). The fitted width of *B* is 33.6 *μ*m. (d) The profiles of *C* and *B* at *T* = 0.01 s for the same parameters as in (a) but with *B*_tot_ = 16, 000 *μ*M. The fitted width of *B* is 15.0 *μ*m.

#### a. Transverse deformation and twisting during contraction

We first consider additional deformation modes besides uniaxial compression. For instance, during contraction of *Spirostomum* there is appreciable twisting of the organism about its compression axis as well as heterogeneous variations in its transverse width. In *Vorticella*, the contraction force is localized to a thin myoneme fiber which wraps around a thicker elastic stalk. Contraction of the approximately one-dimensional myoneme fiber significantly deforms the stalk, pulling it from an initially straight conformation into a compressed helical conformation. A numerical model of this process based on the Cosserat rod theory has been studied in Ref. 48. Interestingly, it was found that the elasticity and dynamics of the stalk only marginally affects the projected one-dimensional compression dynamics, which are to a good approximation dominated by the drag on the large head at the end of the stalk.

Accurate modeling of non-compressional degrees of freedom depends on the specific myoneme arrangement and body plan of the organism under consideration, making it difficult to offer a general prescription for treating these effects. However, accounting for such modes is expected to improve quantitative accuracy since these modes can introduce timescales which may enter in the compression dynamics, depending on the parameter regime under consideration. Additionally, understanding the slow, active re-extension phase of these organisms likely requires detailed modeling of their geometry (see Ref. 48 for a study of re-extension in *Vorticella*). Here, we focus on one deformation mode, which is the twisting during compression experienced by *Spirostomum*. We leave detailed studies of other geometrical aspects of myoneme contraction to future work, although in Section B 3 c we discuss how some of these effects could be approximately treated as passive auxiliary springs which resist the active compression of myonemes.

**FIG. 16.**
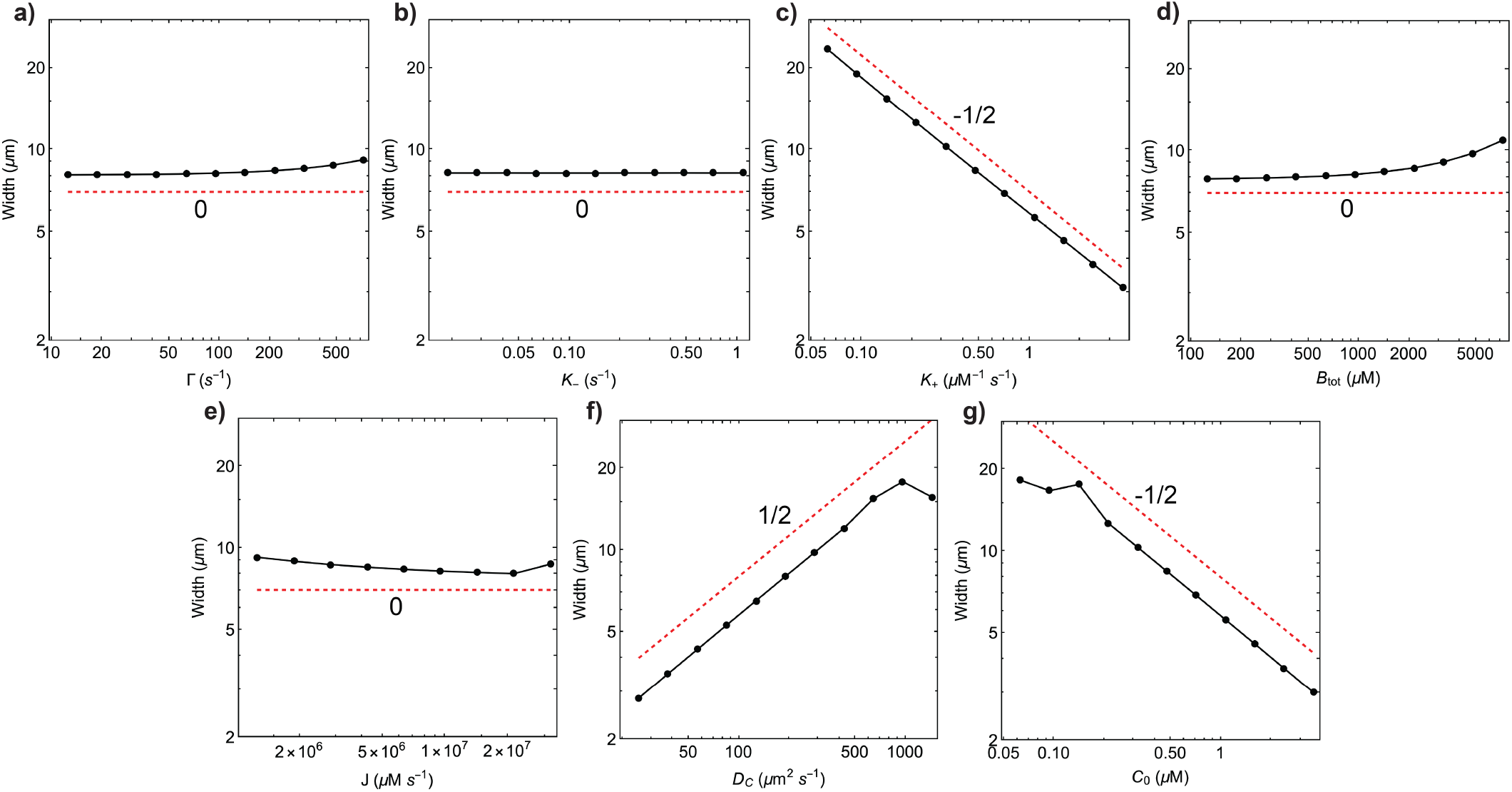
The scaling of *W*. Results from fitting Eq. (B32) to *B*(*S*, 0.005 s), obtained from numerical integration, shown as connected black dots. The red dotted lines illustrate the predicted scaling from the analytical result 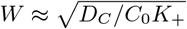.

In *Spirostomum*, a sheath of microtubules wraps around the organism^11,35^. We can view this sheath as a torsional elastic spring which resists twisting and compression. We introduce a variable Θ(*S, T*) describing the local angle that the cross-section makes with the compression axis. A quadratic elastic energy function for the *i*^th^ connection the discrete chain can be written as

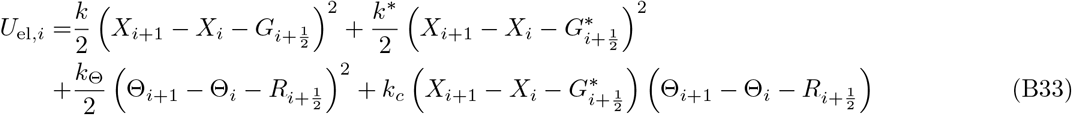

where 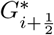 is the rest length of the microtubule sheath segment, 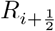 is the equilibrium value of the twist, and *k*_*c*_ captures an energetic coupling between twisting and stretching. We note that positivity of the energy requires setting 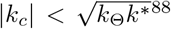 ^88^. This form for the energy is commonly used to model molecules such as DNA^58^. Through steps analogous to those in the previous section, one can pass to a continuum limit of the dynamics of Eq. (B33). Two new timescales are introduced, corresponding to the parameters *k*_Θ_ and *k*_*c*_. We leave a study of this model to future work, but we note here that the model without twisting can be obtained as a limit of Eq. (B33) in which the coupling constant *k*_*c*_ is negligible. The magnitude of this coupling term tends to be much smaller than the stretching modulus (by a ratio of ∼ 1*/*10 for DNA^58^) and can be omitted in the minimal model studied in this paper.

#### b. Poroelasticity

Beyond twisting, an additional dynamical feature of *Spirostomum* contraction is the motion of the internal of fluid, whose possible volume conservation in the organism may produce motion transverse to the contraction direction (see Figure 9 and Ref. 35). Accurate treatment of this aspect of the dynamics requires specifying the poroelastodynamics of the fluid in the cell interior and boundary conditions that prevent its exit through the cell membrane (see, e.g., Ref. 85). While a full treatment of these dynamics is beyond the present study, we now describe a treatment of poroelasticity that provides an approximate understanding of the contribution of the fluid motion to the one-dimensional compression dynamics.

##### a. Poroelastodynamic theory

Poroelasticity describes the mechanics of sponge-like materials that have a connected solid skeleton with pores that are filled with a fluid. Originally developed to describe the mechanics of water saturated soil^89,90^, it is known to introduce timescales in the dynamics of intracellular environments^60,86^, nastic plants^91,92^, and muscles^85^. To treat the effects of poroelasticity on the contraction dynamics of myoneme networks, we build on the linear poroelastodynamic theory outlined in Refs. 93 and 94.

**FIG. 17.**
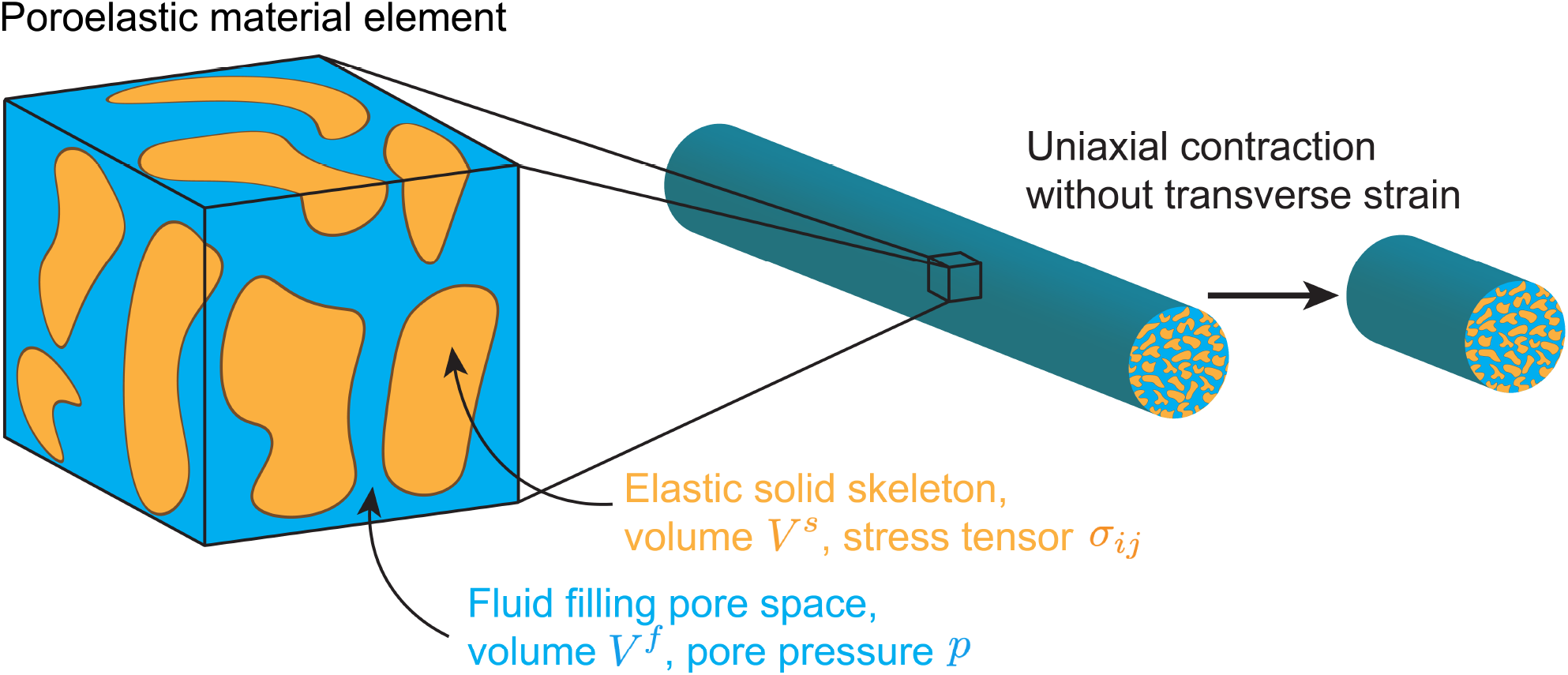
Poroelasticity. Schematic diagram of a poroelastic volume element (left), as well as an illustration of uniaxial compression without transverse strain (right).

A volume element of a poroelastic material has fluid volume *V* ^*f*^ and solid volume *V* ^*s*^, and these together define the porosity

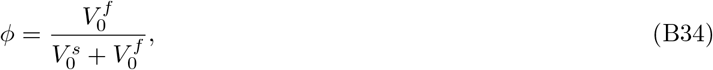

where the subscripts indicate their values in the initial undeformed configuration. For a homogeneous system *ϕ* is constant in space, and because it refers to the undeformed configuration it is also constant in time. The dynamical variables include the displacement of the solid skeleton 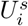 and the displacement of the fluid 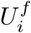, from which we define the specific discharge of the fluid relative to the skeleton 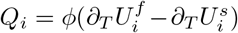. Here, the index *i* = 1, 2, 3 refers to the three Cartesian coordinates. From the motion of the solid skeleton, the symmetric strain tensor 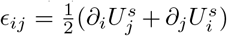) is formed. One also tracks *ζ*, the variation in fluid volume per unit reference volume.

The internal energy of the poroelastic material can be written as a linear function of the kinematic “strain” variables *E*_*ij*_ and *ζ* multiplied by their conjugate “stress” variables *σ*_*ij*_ and *p*. The fluid pore pressure *p* is the interpreted as the pressure of a hypothetical fluid reservoir in equilibrium with the immersed material element. The constitutive equation for the stress tensor *σ*_*ij*_ of an isotropic poroelastic material can be written as^89,90,93,94^

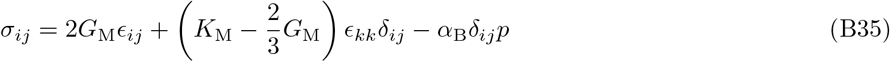

where *G*_M_ is the shear modulus, *K*_M_ is the bulk modulus, *δ*_*ij*_ is the Kronecker delta, and *α*_B_ is Biot’s effective stress constant, a parameter in the range [0, 1] which captures the effect of the pore pressure on the elastic strain energy of the solid skeleton. Repeated indices imply summation, so that *E*_*kk*_ is the trace of the solid skeleton strain tensor. The constitutive equation for the pore pressure is^89,90,93,94^

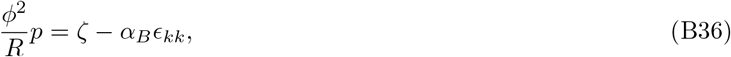

where *R* is a material parameter with units of an elastic modulus. In Ref. 95, Eq. (B36) is derived by accounting for the various factors affecting the proportional change *ζ* of fluid volume in a material element, and it reflects the fact that because *p* is a pressure, it can only couple to the isotropic strain quantities *E*_*kk*_ and *ζ* in the linear constitutive equations^89^. See Ref. 96 for an interpretation of the parameters *α*_B_ and *R* in terms of microscopic properties of the material.

To study the dynamics of a poroelastic material, we next introduce the inertial force balance condition^93,94^

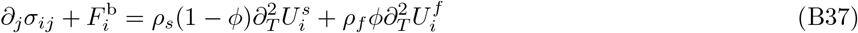

where 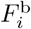 is a body force density, *ρ*_*s*_ is the density of the solid skeleton, and *ρ*_*f*_ is that of the fluid. Local continuity of the fluid also requires that^93,94^

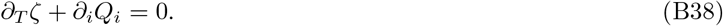

These dynamical equations are closed using the dynamic version of Darcy’s law, which reads^93,94^

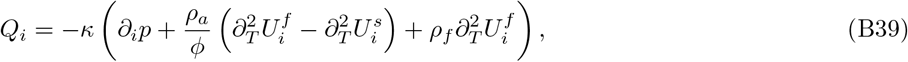

where *κ* is a permeability constant that depends on the fluid viscosity and structural factors of the skeleton, and *ρ*_*a*_ ∝ *ϕρ*_*f*_ is the apparent mass density.

##### b. Extensions to incorporate the physics of myoneme networks

Eqs. (B35) to (B39) represent a closed system describing the poroelastodynamics of a material, and they can be rearranged as differential equations in the variables 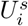 and *p* (although other pairs of variables may be used in equivalent formulations). We now consider two extensions to these equations which represent additional physics in the myoneme-based system.

First, we allow for a non-autonomous active stretch tensor *g*_*ij*_ which depends explicitly on space and time. It is also symmetric, i.e., *g*_*ij*_ = *g*_*ji*_. The elastic restoring force of the solid skeleton should only depend on the relative strain tensor *ϵ*_*ij*_ −*g*_*ij*_, so we use this in place of the absolute strain tensor *ϵ*_*ij*_ where it appears in Eq. (B35), which is now written as

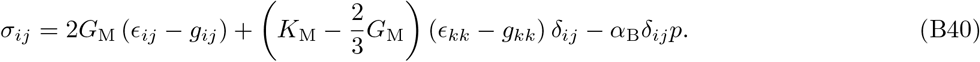

As discussed in Ref. 95 where a similar “thermal strain” is considered in the context of soil dynamics, the constitutive equation for *p*, Eq. (B36), is augmented to account for the variation in fluid volume due to the change in effective stress caused by the non-autonomous active stretch. We thus update Eq. (B36) to

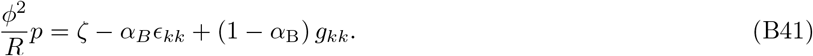

Note that we are using displacement variables *U*_*i*_ here, rather the position variables *X*_*i*_ which are used in the main text. The two formulations are equivalent since the position variables can be recovered by adding the initial positions 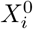 to the displacement variables. If the initial configuration defines the material coordinates, i.e., 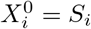, one can show that the interpretation of *g*_*ij*_ here is the same as that in the main text apart from an added constant term 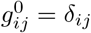. Because derivatives of *g*_*ij*_ and *U*_*i*_ below enter in Eq. (B43) and its approximations below, the constant term and the distinction between displacement and position can be neglected there.

Second, we allow for an additional external source of Stokes drag to act on the solid skeleton as a body force density 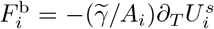 where 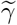 was introduced in *SI Appendix Supplementary methods* Section B 1 and *A*_*i*_ is a cross-sectional area perpendicular to the motion 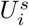. Our simple treatment of the drag as Stokesian and uniform along the system’s length could be improved by introducing a memory kernel for the force history or by letting the drag *μ* be a function of position. However, it was found experimentally that the Stokesian drag dominates the contribution from the memory term in *Vorticella*^41^, and we find here that a uniform *μ* still allows for excellent fits to experimental traces of length during contraction. The explicit dependence of drag on the organisms’ shapes (i.e., cigar-shaped for *Spirostomum* and wine glass-shaped for *Vorticella*) could also be considered in future refinements of the model.

##### c. Simplifying assumptions

The previous equations are not analytically tractable, so we now make two key approximations. First, we assume that the motion occurs only in the *x*-direction and does not produce transverse strain in the *y* or *z* directions. We let the cross-sectional area in the *yz* plane be *A*, which we assume to be independent of *x*, and we denote the material coordinate in the *x* direction as *S*. This motion, visualized in Figure 17, allows us to set *ϵ*_11_ = *∂*_*S*_*U*^*s*^ (where we dropped the subscript on 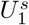) and *g*_11_ = *g* and all other elements of *ϵ*_*ij*_ and *g*_*ij*_ to zero. Second, we assume that the static version of Darcy’s law,

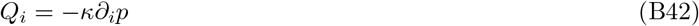

can be used in place of the dynamic version in Eq. (B39). The accuracy of this approximation improves if *κ* is small, as discussed in Refs. 94 and 95.

Under these two approximations, we can write the dynamical equations for the displacement of the solid skeleton in the *x*-direction *U*^*s*^(*S, T*) and the pore pressure *p*(*S, T*) as

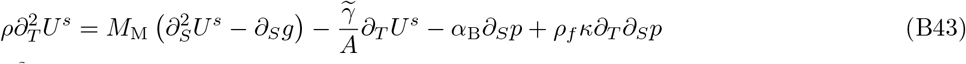

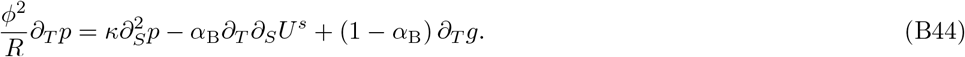

where *M*_M_ = *K*_M_ + 4*G*_M_*/*3 is the p-wave modulus and *ρ* = (1 −*ϕ*)*ρ*_*s*_ + *ϕρ*_*f*_ is the bulk density. For a material whose Poisson ratio is zero, the p-wave modulus describing constrained uniaxial deformation is equal to the Young’s modulus *E*_M_ for unconstrained uniaxial deformation, which for simplicity we assume holds here. In other words, we set *M*_M_ = *E*_M_.

Eqs. (B43) and (B44) should be compared to Eq. (B4) to understand the new physical ingredients due to poroelasticity. To put them in the form of Eq. (B5), we write these new equations as

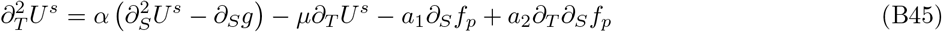

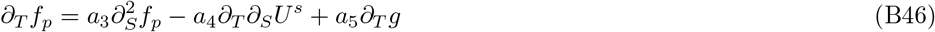

where *f*_*p*_ = *pA* is the force in the *x* direction obtained by multiplying the pore pressure by the cross-sectional area *A*. We have also introduced the constants

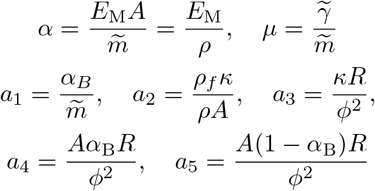

where 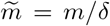 was introduced previously as the linear mass density of the spring chain. This is related to the volumetric bulk density through 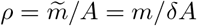.

##### d. Analysis of poroelastic dynamics

Rather than attempt a bottom-up parameterization of the system to fix the seven parameters, we make one further approximation by assuming that both the fluid and solid skeleton are incompressible. This is known as the “incompressible constituents” assumption, and its implications in the context of cytoplasmic poroelasticity of eukaryotic cells (including its association with pressure diffusion dynamics) is described in Ref. 60. As detailed in Ref. 94, this assumption corresponds to *α*_B_ = 1, and we thus set *a*_5_ = 0. It further corresponds to *R*→ ∞ (which can be shown to not affect setting *a*_5_ = 0). This implies that the large terms proportional to *a*_3_ and *a*_4_ on the right-hand-side of Eq. (B46) cause the poroelastic force *f*_*p*_ to equilibrate rapidly with respect to the change in *U*^*s*^. This is a quasi-steady state limit^97^ for the dynamics of *f*_*p*_, allowing us to set *∂*_*T*_ *f*_*p*_ = 0. Eq. (B46) then implies that

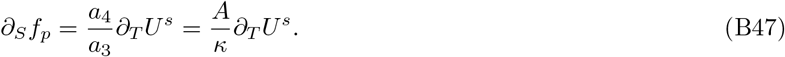

Substituting this relation in Eq. (B45) and setting the term proportional to *∂*_*T*_ *∂*_*S*_*f*_*p*_ to zero, we have

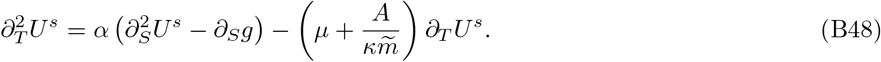

In other words, for this limit the effect of poroelasticity is simply to add a term to the Stokesian drag on the motion of the solid skeleton. Under this limit, the scaling results obtained in the main text using the model without poroelasiticty still hold because the effective drag 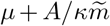 is a linear function of the external drag *μ*, having a constant additional contribution 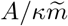 which is determined by the permeability, density, and size of the organism’s interior. We leave to future work a numerical exploration of Eqs. (B45) and (B46), where the incompressible constituents approximation is not made. Measuring the pororoelastic parameters for *Spirostomum* and considering models that account for transverse strain (which can be appreciable, as shown in Figure 9) are additional important directions for future work.

#### c. Auxiliary elastic elements

Finally, we extend the model by including an auxiliary elastic element which passively resists compression caused by the Ca^2+^-activated springs. This represents a simplified model for, for instance, the effect of the microtubule sheath in *Spirostomum* or the bent elastic stalk in *Vorticella*^35,48^.

In the discrete model, the force on the *i*^*th*^ node becomes

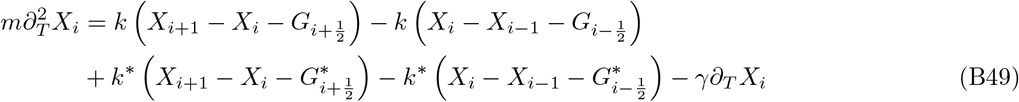

where the starred parameters refer to a set of auxiliary springs acting in parallel to the active springs. We assume that the auxiliary spring rest lengths *G*^*\**^ are uniform and constant in time. Passing to the continuum limit, we have

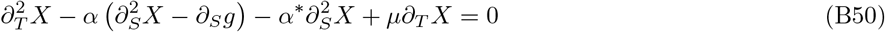

which can be re-expressed as

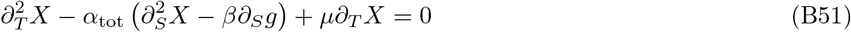

where *α*_tot_ = *α* + *α*^*\**^ and *β* = *α/α*_tot_ ∈[0, 1]. The force balance at the end of the chain imposes the boundary conditions

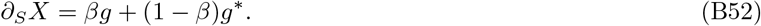

In the steady-state, we have

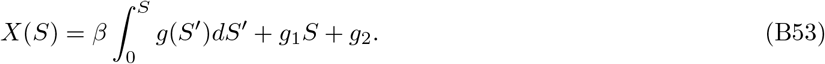

The integration constant *g*_2_ can be set to zero, and the boundary conditions imply that *g*_1_ = (1− *β*) *g*^*\**^. At long times, *g*(*S*) = *g*_min_, so that the steady-state is

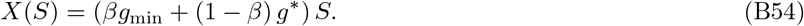

The equilibrium shape is thus determined by a weighted combination of the stretches *g*_min_ and *g*^*\**^ with the weights determined by the corresponding stiffness of the active and auxiliary springs.

We next ask whether the auxiliary springs affect the dynamical scaling results obtained without auxiliary springs, when *α*^*\**^ = 0. By defining 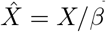, we can map Eq. (B51) into

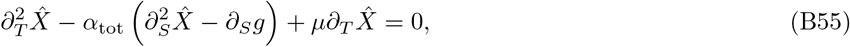

which is formally identical to Eq. (B5). The same logic leading to the scaling relationships for 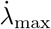 obtained in the main text now carry through for 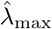, which is defined using 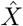 (and 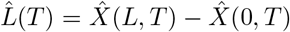) in place of *X*. In the new scaling relations, *α* is replaced by *α*_tot_ and *L* is replaced by 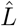.

### 4. Exact solution in the quench limit

The solution to Eq. (15) in the main text can be found exactly using a Fourier expansion. The initial condition is expanded as

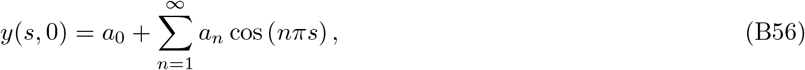

where *a*_0_ may be set to zero and

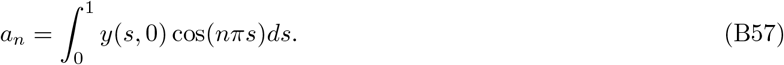

By linearity, the solution can then be written as

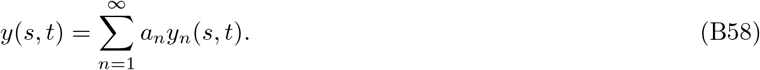

The functions *y*_*n*_(*s, t*) can be found by solving Eq. (15) in the main text with the simplified initial conditions *y*(*s*, 0) = cos(*nπs*), and the result is

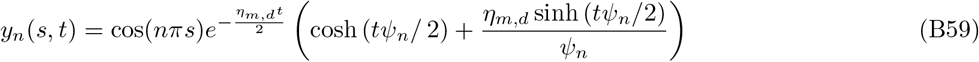

where

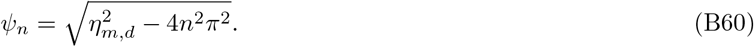

There is a similarity between the continuum quench model, whose dynamics are given in Eq. (15) of the main text, and perhaps the simplest dissipative contraction dynamics which one could consider: the motion of a single mass attached by a spring to a wall, following sudden release from an initial fixed state. The position of the mass *z*(*t*) obeys

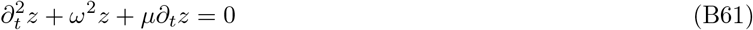

where 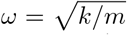 and *μ* = *γ/m* are defined in terms of the spring’s stiffness *k*, mass *m*, and the drag coefficient *γ*. By rewriting *y*_*n*_(*s, t*) from Eq. (B59) as

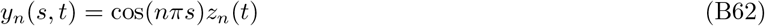

and using the solution of *y*(*s, t*) from Eq. (B58), one can show that Eq. (15) in the main text is equivalent to

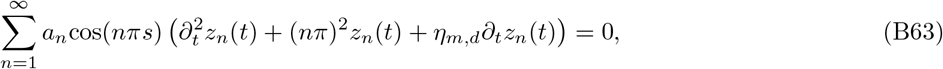

which further implies that

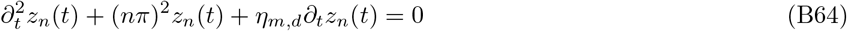

for each *n*. Thus, in the continuum quench model, each spatial harmonic of the initial deviation from equilibrium behaves like a damped spring with frequency *nπ* and drag *η*_*m,d*_.

### 5. Energetics of contraction

Here we consider the energetics of myoneme contraction. The elastic energy of the system is the sum of the energy of each spring in the chain. In the continuum limit this is (cf. Eq. (2))

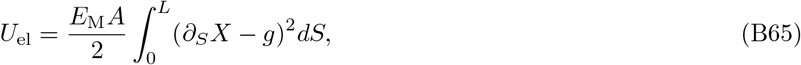

using dimensional variables. The non-dimensional version is

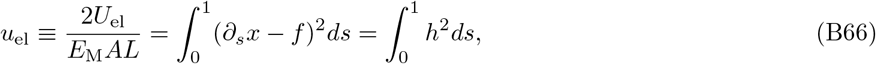

where we have absorbed the factor 1*/*2 into the definition of *u*_el_ for simplicity and used the definition of *h* in Eq. (12) of the main text. Trajectories of *l*(*t*) and *u*_el_(*t*) are shown in *SI Appendix* Figure 7 for several values of the non-dimensional model parameters *η*_*w,m*_ and *η*_*w,d*_.

In the quench limit, the system starts at rest with *x* = *s* and *f* = 1 and all springs then instantly take new rest lengths *g*_min_, causing the internal energy to jump discontinuously from *u*_el_ = 0 to (1 −*g*_min_)^2^. We can regard (1 −*g*_min_)^2^ as the maximum internal energy that the system can store in response to a active stretch wave *g*(*s, t*) taking values in [*g*_min_, 1]. Outside the quench limit, the system relaxes while the wave is traversing its length, and the maximum internal energy stored in the organism during the contraction trajectory 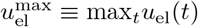 is less than (1 − *g*_min_)^2^.

We can therefore quantify how quench-like the dynamics are using

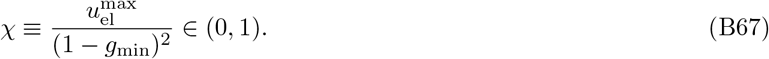

